# Precision Proteoform Design for 4R Tau Isoform Selective Templated Aggregation

**DOI:** 10.1101/2023.08.31.555649

**Authors:** Andrew P. Longhini, Austin DuBose, Samuel Lobo, Vishnu Vijayan, Yeran Bai, Erica Keane Rivera, Julia Sala-Jarque, Arina Nikitina, Daniel C. Carrettiero, Matthew Unger, Olivia Sclafani, Valerie Fu, Michael Vigers, Luc Buee, Isabelle Landrieu, Scott Shell, Joan E. Shea, Songi Han, Kenneth S. Kosik

## Abstract

Prion-like spread of disease-specific tau conformers is a hallmark of all tauopathies. A 19-residue probe peptide containing a P301L mutation and spanning the R2/R3 splice junction of tau, folds and stacks into seeding-competent fibrils and induces aggregation of 4R, but not 3R tau. These tau peptide fibrils propagate aggregated intracellular tau over multiple generations, have a high β- sheet content, a colocalized lipid signal, and adopt a well-defined U-shaped fold found in 4R tauopathy brain-derived fibrils. Fully atomistic replica exchange molecular dynamics (MD) simulations were used to compute the free energy landscapes of the conformational ensemble of the peptide monomers. These identified an aggregation-prohibiting β-hairpin structure and an aggregation-competent U-fold unique to 4R tauopathy fibrils. Guided by MD simulations, we identified that the N-terminal-flanking residues to PHF6, which slightly vary between 4R and 3R isoforms, modulate seeding. Strikingly, when a single amino acid switch at position 305 replaced the serine of 4R tau with a lysine from the corresponding position in the first repeat of 3R tau, the seeding induced by the 19-residue peptide was markedly reduced. Conversely, a 4R tau mimic with three repeats, prepared by replacing those amino acids in the first repeat with those amino acids uniquely present in the second repeat, recovered aggregation when exposed to the 19- residue peptide. These peptide fibrils function as partial prions to recruit naïve 4R tau—ten times the length of the peptide—and serve as a critical template for 4R tauopathy propagation. These results hint at opportunities for tau isoform-specific therapeutic interventions.

**Significance Statement:** A structural motif corresponding to a short junction sequence spanning R2 and R3 forms fibrils that adopt a fold characteristic of 4R tauopathy fibrils and induces misfolding of the larger tau protein with loss of microtubule binding and a prion-like specificity for 4R tau. Simulations, validated experimentally, pinpointed the specific amino acids in the peptide that can toggle its properties between aggregation competent and incompetent. The modifications suggest design principles for a therapeutic intervention potentially capable of disaggregating tau or preventing its aggregation in the 4R tauopathies.

## Introduction

The pathogenetic mechanisms of tauopathies critically involve a prion-like spread of misfolded tau, often displaying disease-specific topographies and conformations (1–3). Prion-like tau protein spread can be defined as strain-specific generation and propagation of a distinct amyloid shape within the brain (4–6). As tau strains spread from cell to cell, the donor protein accurately templates the misfolding of tau in the recipient cells. Capturing tau for templating requires the prion-like donor tau to select a tau conformer from a broad energy landscape containing an ensemble of conformers that will lead to the replication of misfolded tau with the specific atomic level structures as adopted in fibrils diagnostic of the clinical phenotypes of tauopathies, such as chronic traumatic encephalopathy (CTE), frontotemporal dementias, and Alzheimer’s disease (AD) (7). The exact structural and dynamic molecular details of how tau transitions to specific amyloid forms under different clinical circumstances and cellular environments are still not well understood but will be necessary to produce the specific pathological conformations *in vitro*. The intrinsically disordered tau population is in stable equilibrium between monomers, dimers and trimers according to a recent study using single-molecule mass photometry. However, the native-like oligomers present at equilibrium with the monomer, in the absence of aggregation-inducing factors, are thought to be off-pathway with respect to fibrilization according to recent studies (8,9), Instead, the relevant species may be distinct conformers that tau adopts, depending on the energy landscape of the conformational ensemble of tau monomers that promote self-association by stabilizing intermolecular contacts, leading to oligomerization (10), fibrillization, and seeding in recipient cells (11). The companion paper by Vigers et. al. demonstrated that the widely used P301L mutation dramatically alters the energy landscape of the conformational ensemble of tau monomers, and as a result drives up the population of the aggregation-competent conformers.

Two highly studied six amino acid peptides—VQIINK in repeat domain 2 (R2) and VQIVYK in repeat domain 3 (R3)—can assemble *in vitro* into β-sheet-rich fibrils, but these fibrils are not seed-competent (12–14). Characteristics of these sequences known as PHF6* and PHF6, respectively, have been studied through the clearance capacity of peptide mimetics targeting these amyloidogenic motifs (15–17), through nanobodies targeting PHF6 (18), and by enhancing the function of the chaperones Hsp40, Hsp70, and Hsp90, known to bind around PHF6 (19). While these short sequences in tau are recognized as amyloidogenic *in vitro*, little is known about the additional molecular features that confer the property of seeding. Attempts to define a minimal seed have included tau trimers (20), individual repeat peptides (21), a seed-competent monomer (22), peptides that extend the amyloidogenic hexapeptides (12,13,23), a 31-amino-acid peptide (residues 306-336) (23) and a 17-amino-acid peptide (residues 295-311) (23), but our understanding remains incomplete and a rational design of a tau prion with predicted strain specificity has not been demonstrated to date.

The companion paper by Vigers et. al. takes a step toward this so far elusive goal. For the fold adopted by the fibrillar tau to propagate monomeric tau by templating, tau monomers must stack in-register onto fibrils for prion-like propagation. Given that every distinct tau fibril structure solved to date includes PHF6 in the fibril core, it must be included in the fibril interface and stabilize a specific fold. The companion paper highlights that the amyloidogenic PHF6(*) motifs must be oriented such that the backbone hydrogen-bonding moieties that form the intermolecular β-sheets are positioned along the fibril growing axis, and not occupied by intra-molecular β-hairpins. Shape propagation by templated seeding cannot be fulfilled with standalone PHF6 or PHF6* fragments that can freely rotate unless these fragments are stably held in a specific orientation with a counter-strand when forming fibrils. The 19-residue peptide fragment at the R2/R3 junction, jR2R3, exactly achieves that requisite with the PHF6 held in place by a counter-strand stabilized by intramolecular salt-bridge and/or hydrophobic interactions, and the backbone hydrogen-bonds oriented along the fibril growing axis. The counter strand not only stabilizes the aggregation-competent orientation for PHF6, but it also offers additional shape stability. The 3 Å cryo-EM structure of the jR2R3-P301L fibril of the companion paper revealed that this 19-residue peptide adopts the conformation found in 4R tauopathy fibrils at the R2/R3 splice junction, in which a minimal N-terminal segment 295- 300, as part of the second repeat of 4R tau, stabilizes and holds in place the PHF6 segment within a U-shaped fold (PDB: 8V1N). The fibrils formed of jR2R3-P301L, with four peptide chains folded and packed in 2D across the fibril cross section, are highly homogeneous, abundant, stable, and seeding competent to recruit 4R tau *in vitro*. However, for jR2R3-P301L fibrils to qualify as tau prion, they must misfold naïve tau intracellularly, propagate aggregation in a strain or isoform-specific manner, and generate insoluble tau that maintains such activity.

This study presents the design of a small tau peptide with prion-like properties that detail the faithful templating of 4R tauopathies. These peptide-induced 4R tau aggregates are sarkosyl-insoluble, high molecular weight assemblies that robustly propagated to daughter cells across multiple generations and passages and contain a lipid signal localized to the aggregate. We simulated the detailed molecular dynamic interaction of a seed with a localized area of native tau and validated these predictions experimentally. Experiments revealed a mechanism that lowered the energy barrier toward 4R tau oligomerization by breaking intramolecular hydrogen bonds and exposing the amyloidogenic PHF6 region such that its hydrogen bond forming functionalities were unoccupied and available for recruiting tau monomers to the active fibril end. Importantly, these observations suggest a design principle for preventing aggregate formation. Re-positioning very few of the residues that define the 4R tau as in or out of the sequence could control aggregate formation. Of particular interest was the single S305K substitution in cellular tau that prevented seeding with the otherwise 4R specific seeding competent peptide. This mutation that rendered cellular 4R tau as 3R tau-like at one site pinpointed a structural feature involved in templating. A search among the tau fragments known to exist in tauopathy brain tissue (25–27) for those that resemble the small probe peptide used here could reveal a more precise therapeutic target.

## Results

### A Small Tau Peptide Induces Tau Fibrils

Two 19-amino-acid tau peptides (295–313) designated jR2R3 and jR2R3 P301L, that differ only by the presence of the P301L mutation in jR2R3 P301L (Figure 1A), can readily form fibrils (23). jR2R3 and jR2R3 P301L span the splice junction between the second and third repeat and contain the PHF6 aggregation motif (VQIVYK) (12, 15, 18). PHF6 appears in various contexts within the recently solved full-length tau fibril structures, always adopting β-strand conformations (29–32). For tauopathies containing exclusively 4R tau (corticobasal degeneration (CBD), argyrophilic grain disease (AGD), aging-related tau astrogliopathy (ARTAG), progressive supranuclear palsy (PSP), globular glial tauopathy (GGT), GGT-PSP tauopathy (GPT)), the jR2R3 sequence folds into a strand-loop-strand motif (Figure 1B) (33). In pure 3R tauopathies, such as Picks’ disease (PiD), which lack residues present in jR2R3 and jR2R3 P301L, the PHF6 amyloidogenic sequence does not reside in the context of a strand-loop-strand (33). For tauopathies with mixed 3R/4R isoforms (Alzheimer’s disease (AD), Chronic Traumatic Encephalopathy (CTE)), the N-terminal half of the jR2R3 sequence—residues 295-305—is not present in the solved Cryo-EM structures(29, 30) (Figure 1B), but NMR studies show that 3R and 4R tau are well mixed in AD-tau seeded fibrils, and the mixed fibrils reveal no structural differences in the rigid β-sheet core or the mobile domains(35). These structural data are important for understanding the prion-like properties of the jR2R3 peptides and suggest that templating and propagation are driven by different molecular mechanisms and structural motifs in different tauopathies.

**Figure 1:**
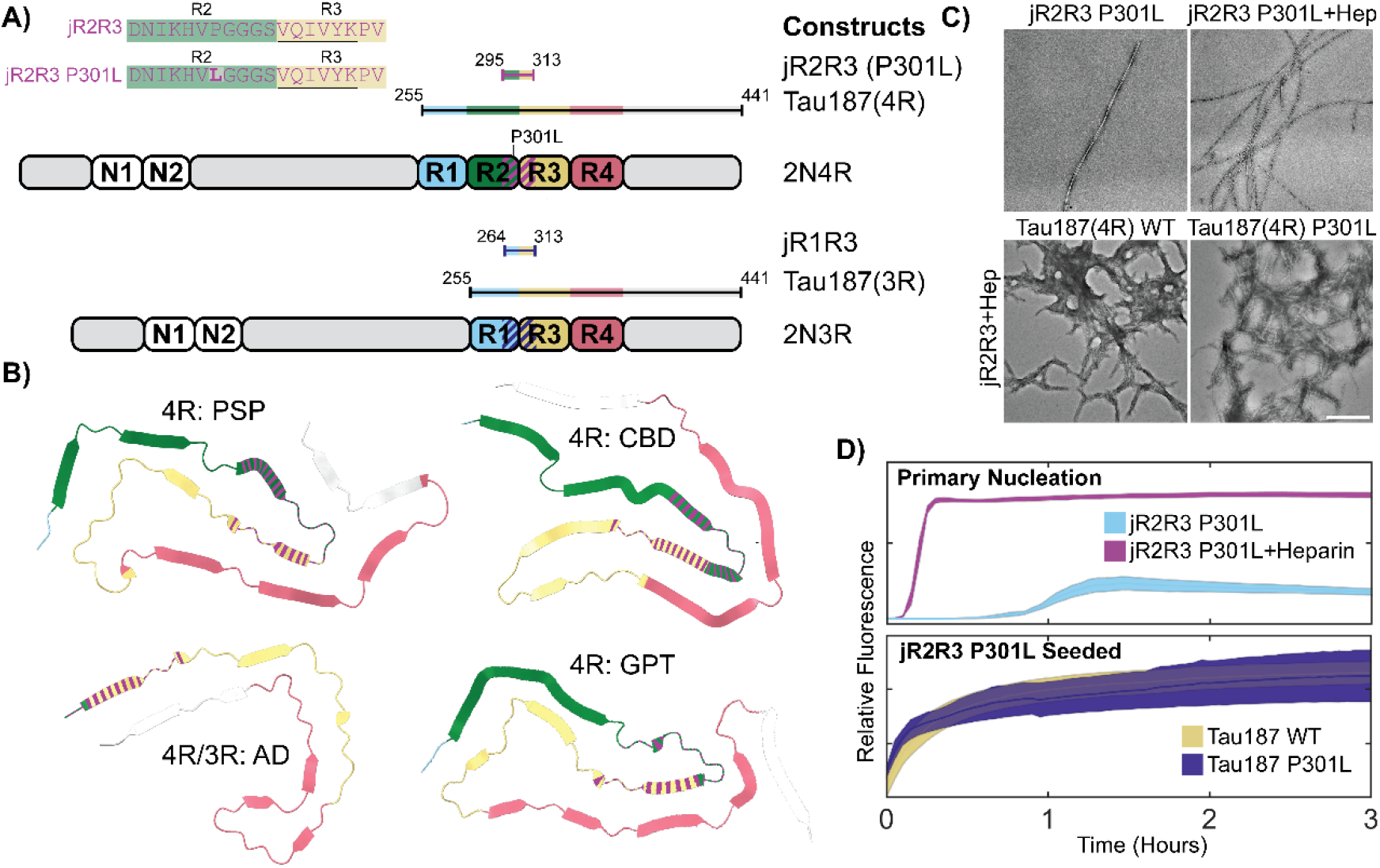
Tau Constructs, Structural Representations, and Fibrilization Characteristics of jR2R3 and jR2R3 P301L. A) Overview of the tau constructs utilized in the study, including 3R and 4R isoforms. These strand loop strand sequences, designated jR2R3 and jR2R3 P301L, are shown in the top left with the PHF6 sequence underlined and residues mutated relative to jR2R3 shown in bold. The diagram also illustrates the most commonly used constructs, categorizing jR2R3, jR2R3 P301L, and Tau187(4R) as 4R tau constructs, and categorizing Tau187(3R) and jR1R3 as 3R tau constructs. The elements are color-coded for clarity with R1 highlighted in Cyan, R2 in Green, R3 in Sand, R4 in Rose, and jR2R3 and jR2R3 P301L in Purple. Dashed line on 2N4R and 2N3R represent the jR2R3/jR2R3 P301L junctions. B) PDB structures representing 4R tauopathies (PSP with PDB ID: 7P65(32), CBD with PDB ID: 6VHA(66), GPT with PDB ID: 7P6A(32), and mixed 4R/3R tauopathy (AD with PDB ID: 6VHL(66). The jR2R3 and jR2R3 P301L constructs are centrally located in 4R tauopathies and are hypothesized to be minimal elements capable of specifically templating 4R isoforms. In mixed 4R/3R and 3R tauopathies, jR2R3 and jR2R3 P301L are absent due to splicing of the R2 repeat, which would hinder templating by jR2R3 and jR2R3 P301L due to clashes with the R1 alternatively spliced sequences in 3R sequences. The structures are colored with the conventions established in (A), with jR2R3 and jR2R3 P301L represented by striped coloration over their respective sequences. C) TEM images displaying the fibrilization of jR2R3 P301L in the absence (Top Left) and presence of heparin (Top Right), as well as seeding of jR2R3 P301L with heparin seeds. Scale Bar: 500 nm. D) (Top) ThT assays comparing jR2R3 P301L fibrilization alone (cyan) and with heparin (purple). (Bottom) ThT assays after adding jR2R3 P301L heparin-seeded fibrils to Tau187 WT (sand) and Tau187 P301L (indigo) monomers. Error bounds represent standard error of the mean.

jR2R3 P301L demonstrated the capacity to fibrilize under a wide range of conditions (Figure 1C, 1D and Vigers et. al., companion paper). In contrast, jR2R3 could only aggregate in the presence of heparin and shaking, and even under these conditions, only some of the reactions resulted in productive fibril formation, which took more than 24 hours (Figure S1A, S1B). The aggregation of jR2R3 P301L, with and without heparin, was rapid, reaching plateaus in less than three hours. The companion paper demonstrated that the U-shaped fold of jR2R3 P301L stacks as a strand-loop-strand β-sheet with a hyperlocalized hotspot around sites 300 and 301 capable of promoting in-register pinning of tau for templated aggregation. These aggregates displayed fibril-like morphologies with characteristic periodicity as observed by TEM (Figure 1C).

Next, we tested whether jR2R3 and jR2R3 P301L fibrils could seed native jR2R3 and jR2R3 P301L monomers. All peptide seeds had excess heparin removed to prevent residual cofactor-induced fibrilization. In both cases, jR2R3 and jR2R3 P301L fibrils were able to propagate fibrils from jR2R3 and jR2R3 P301L monomers (Figure S1C, S1D).

We tested if jR2R3 and jR2R3 P301L fibrils could seed the larger tau constructs, Tau187- WT and Tau187-P301L (a.a. 255-441 includes core structures observed in cryo-EM tauopathies) (Figure 1A). Fibrils formed from either of the peptides robustly seeded fibril growth (Figure 1C Bottom, 1D Bottom, S1E, S1F). Although jR2R3 initially has a significant barrier to aggregating into β-sheets, once it overcame this kinetic barrier and formed fibrils, its ability to template new tau monomers for further fibril propagation remained effective *in vitro*.

These seeded fibrils appeared more amorphous, but periodicity and a fibrillar appearance were readily observed. These results show that these short fragments were sufficient for propagating tau fibrils that extend well beyond the core amyloidogenic sequence which directly interacts with the peptide.

### Seeding and Aggregation of jR2R3 P301L Induced Tau Aggregates in Cells

We evaluated seeding in H4 cells stably expressing mClover3-Tau187-P301L, a construct that encompasses the core of all disease tau fibrils solved by Cryo-EM (29–32, 34). jR2R3 P301L fibrils seeded tau aggregates (Figure 2A, 2B Top). At concentrations of 2 μM jR2R3 P301L fibrils, 30 ± 7% of cells exhibited at least one visible tau aggregate eight hours post jR2R3 P301L seeding. The experiments described next were conducted relative to this benchmark: a seeding assay displaying the most cells with puncta (Figure 2A). Full-length mClover3-Tau constructs were evaluated for their aggregation tendencies in the presence of jR2R3 P301L fibrils. Both mClover3-0N4R-P301L, and mClover3-2N4R-P301L robustly formed aggregates when seeded by jR2R3 P301L (Figure 2B, S2A, S2B). mClover3-Tau187-WT and mClover3- 0N4R-WT showed significantly less aggregation than their corresponding P301L variant counterparts (Figure 2B). Additionally, mClover3-2N4R-WT showed less aggregation than the Tau-187 constructs, potentially due to the aggregation inhibitory role of the N-terminus (36) (Figure 2B, S2B Top). More aggregates were observed in cells with mClover3-P301L constructs than the corresponding WT lines for all constructs tested, and Tau187 lines showed more aggregates when compared to full-length tau (Figure 2B Bottom, S2A, S2B). The less well- fibrillized jR2R3 peptides were unable to seed tau aggregation across a range of concentrations up to 5 μM fibril and tested within the first 24 hours after seeding. Neither monomeric jR2R3 nor jR2R3 P301L could seed aggregates in cells at any tested concentration or time scale.

**Figure 2:**
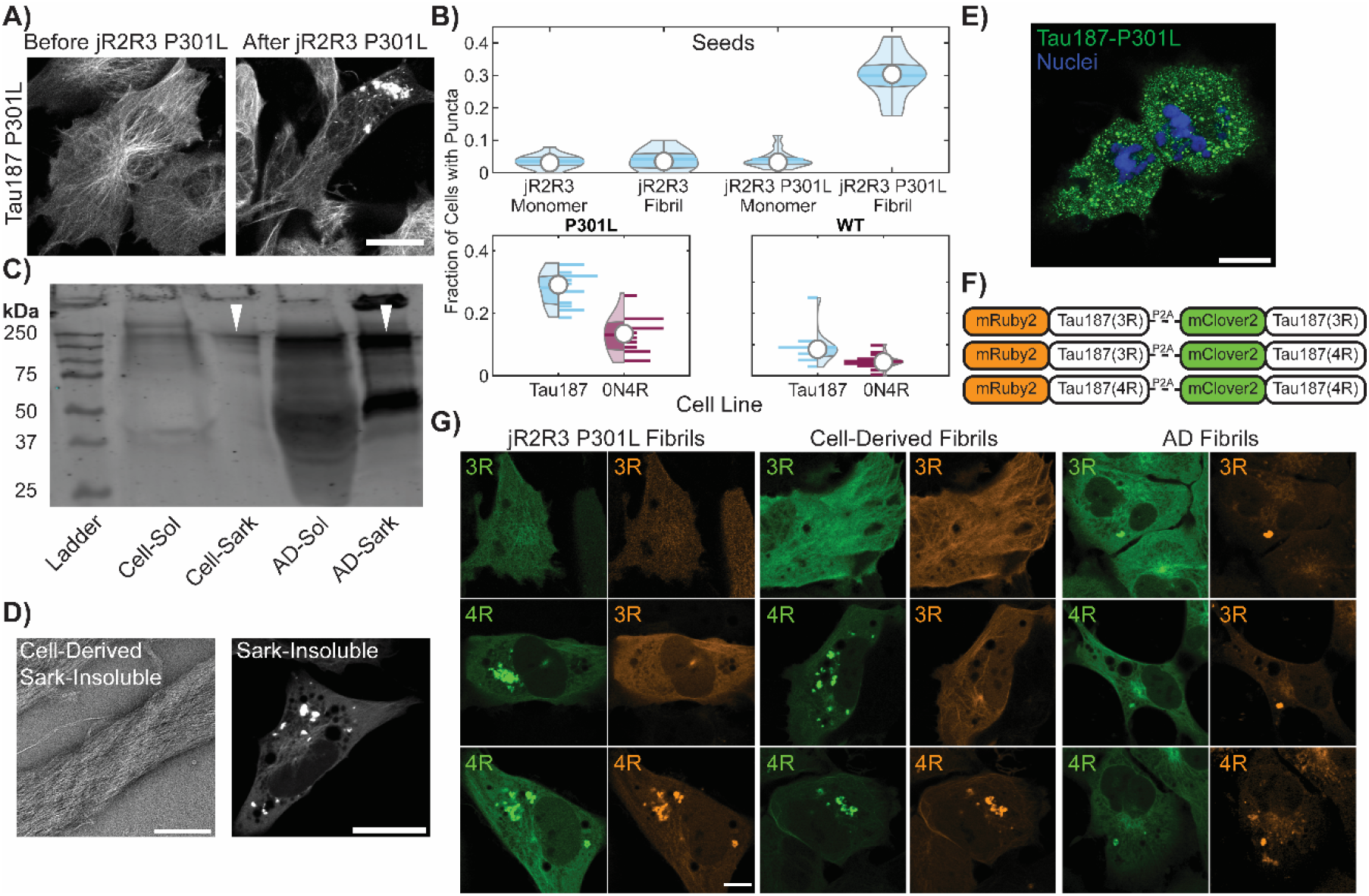
Tau Aggregation Formation, Propagation, and their Characteristics in Cells Seeded with jR2R3 P301L Fibrils. A) H4 cell stably expressing mClover3-Tau187-P301L, seeded with jR2R3 P301L fibrils, imaged at 0- and 12-hours post jR2R3 P301L fibril addition. Scale bar: 20 μm. B) Violin plots of mClover2-Tau187-P301L cell lines seeded with the peptide series of monomers and fibrils (Top). Further analysis focused on jR2R3 P301L fibrils added to cells lines expressing Tau187 P301L or full length 0N4R P301L (Bottom Left) or Tau187 WT and full length 0N4R WT (Bottom right). Percentage of cells with puncta were 5 ± 2.3% of 0N4R WT cells had puncta and 13 ± 6 % of 0N4R P301L cells had puncta. Tau187 lines had the most puncta with Tau187 WT having 9 ± 5.5 % and Tau187 P301L having 28 ± 5 %. C) Western blot of sarkosyl soluble and insoluble material from mClover3-Tau187-P301L cells seeded with jR2R3 P301L and material derived from AD brain samples. Blot stained with the disease-specific conformational tau antibody (MC1). Similar-sized oligomers observed for both cell-derived and patient-derived material highlighted with white triangles. D) (Left) TEM of sarkosyl insoluble material derived from cell lines. Scale bar: 250 nm. (Right) Sarkosyl insoluble fractions were added to the original mClover3-Tau187-P301L cell lines showing seeding of puncta after 12 hours. Scale bar: 20 μm. E) Cells seeded with jR2R3 P301L undergo division and propagate aggregates to daughter cells. Green: Tau187-P301L, Blue: Hoescht. Scale bar: 20 μm. F) Schematic of the three constructs used for testing the selective templating of either the 4R or 3R isoform of tau by jR2R3 P301L. Each construct consists of either the 4R or 3R isoform of tau fused to mRuby3, and is expressed concurrently with another 4R or 3R isoform of tau fused to mClover3. A P2A cleavage site between the two tau fusions ensures equimolar expression of the two fusions from the same expression cassette. G) 12 hours after the addition of jR2R3 P301L, cells expressing two copies of 4R tau showed aggregates in both channels (Left Bottom), cells expressing both 4R and 3R isoforms showed aggregates exclusively in the 4R channel (Left Middle), whereas cells expressing two copies of 3R tau exhibited no aggregates (Left Top). This provides strong evidence that isoform-specific templated elongation is the primary mode of propagation in these cell lines. As a control, fibrils from AD samples, which are characteristic of a mixed 4R/3R tauopathy, seeded both 3R and 4R isoforms as anticipated (Middle). Additionally, cell-derived material was reintroduced to the described cell-lines, and selective 4R seeding was again observed (Right). Scale bar: 20 μm.

Fluorescence recovery after photobleaching (FRAP) experiments indicated that the inclusions did not undergo fluorescence recovery after bleaching. Nine separate mClover3- Tau187-P301L jR2R3 P301L seeded aggregates were completely bleached, and no recovery of fluorescence was observed over several minutes (Figure S3A). Moreover, when large aggregates were half-bleached, there was no redistribution of fluorescence from the unbleached half of the aggregate (Figure S3B). Collectively, these results indicate that the tau inclusions did not display liquid-like exchange properties with the surrounding cellular space, consistent with the expectations that these are solid or gel-like aggregates.

### Extraction and Passaging of Cell Culture Seeded Material

Large quantities of cells expressing mClover3-Tau187-P301L were seeded with jR2R3 P301L fibrils, from which we extracted sarkosyl soluble and insoluble material. Western blots of the soluble material revealed multiple bands labeled with MC1 at ∼240 kDa which would correspond to ∼6-mers of mClover3-Tau187. We then compared these patterns to those from sarkosyl soluble and insoluble fractions of tau extracted from Alzheimer brain samples. MC1 positive oligomer bands ran at molecular weights that approximated 6-mers of full-length tau (Figure 2C). Additionally, a fraction of the insoluble material ran at extremely high molecular weights that did not enter the wells. TEM images of the insoluble material extracted from seeded cells exhibited distinct fibril-like structures (Figure 2D Left). Next, the insoluble material from the cell culture lines was added back to cells expressing mClover3-Tau187-P301L to test its seeding efficacy. The fibril fractions robustly seeded tau aggregation, whereas the soluble fractions did not (Figure 2D Right).

Encouraged by the strain propagation observed with cell-extracted seeding, we cultured jR2R3 P301L seeded cells for extended periods to look for the propagation of aggregates in dividing cells. Cell lines harboring aggregates induced by jR2R3 P301L were passaged at low densities, and their division was monitored. Cells with aggregates underwent cell division, and the resulting daughter cells continued to have aggregates, suggesting authentic propagation of a strain (Figure 2E). By passaging single cells into wells, we could clonally isolate cell lines that propagated the strain specifically over multiple passages.

### Demonstrating 4R Selectivity of jR2R3 P301L Seeding in Cell Culture

If jR2R3 P301L is functioning as a prion-like moiety, we should be able to detect precise templating and propagation of a strain. In 4R tauopathies, 4R is selectively incorporated into fibrils (24, 36) and the fibril structure corresponds to the specific 4R tauopathy, which spreads with high structural fidelity. Considering the complementarity of the jR2R3 and jR2R3 P301L sequences with 4R tau but not 3R tau, along with the cryo-EM structure from our companion paper of jR2R3 P301L fibrils that adopt a core 4R tauopathy fold (24), we hypothesized that jR2R3 P301L should be able to elongate only 4R tau isoforms. To examine selective 4R aggregate formation, we created three cell lines expressing equimolar ratios of 3R and 4R isoforms of Tau187. The two tau constructs, each labeled with a distinct fluorophore (mRuby3 or mClover3), were separated by the self-cleavable P2A sequence. This allowed for visualization isoform-selective incorporation into aggregates (Figure 2F).

As anticipated, when jR2R3 P301L fibrils were introduced into cells expressing two 4R isoforms of Tau187, aggregates were observed in both channels (Figure 2G Left Bottom). In contrast, cells expressing two copies of the 3R isoform of Tau187 showed no formation of aggregates (Figure 2G Left Top). Finally, cells expressing one copy each of 4R and 3R Tau187 only displayed aggregates in the channel corresponding to 4R Tau187 (Figure 2G Left Middle). The cell derived material, sarkosyl extraction of insoluble Tau187 aggregates (after seeding with jR2R3 P301L fibril) was then added to dual expressing 3R/4R Tau187 cell lines (Figure 2G Middle). The same selective seeding of 4R tau, as seen with jR2R3 P301L, was again observed. This outcome demonstrated the 4R selectivity of seeding. In comparison, seeding with AD brain-derived material, induced the formation of aggregates that exhibited fluorescence in both channels in each cell line, consistent with the mixed 3R and 4R tau isoform composition of AD brain fibrils (Figure 2G, right) (37).

### Defining the Energy Barrier to 4R Tau Fibrilization

The observation that jR2R3 P301L could seed Tau187(4R) over Tau187(3R) led us to search for unique features of the R2-R3 junction found in 4R isoforms compared to the R1-R3 junction found in 3R isoforms. Another 19 amino acid peptide that spans the R1-R3 junction is referred to as jR1R3 (Figure 3A). Notably, jR2R3 and jR1R3 differ by only four amino acids while the amyloidogenic PHF6 region is shared in both sequences. We synthesized a peptide corresponding to the jR1R3 sequence and observed that, in contrast to the jR2R3 and jR2R3 P301L peptides, jR1R3 could neither fibrilize on its own, nor in the presence of heparin. None of the conditions tested, including rapid agitation, induced aggregation over any of the timescales tested (data not shown) despite having the amyloidogenic PHF6 sequence. We next added jR2R3 or jR2R3 P301L fibrils to jR1R3 monomers to seed jR1R3 aggregation. Again, no aggregation was observed under any condition (data not shown). Thus, the four amino acid differences between these two peptides conferred strong resistance to fibrillization.

**Figure 3:**
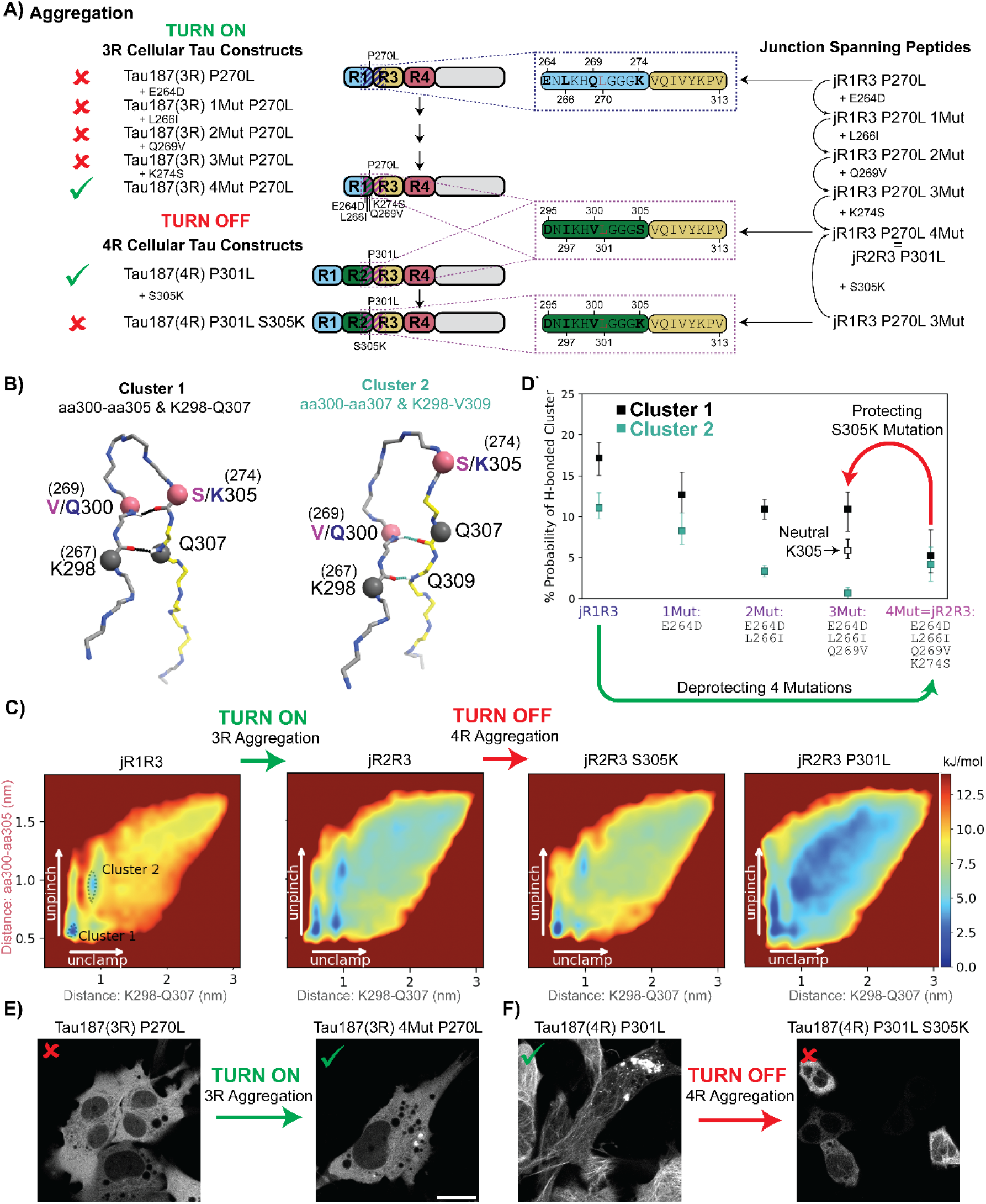
Comparative Analysis of Energy Landscapes and Hydrogen Bonding in jR2R3, jR2R3 P301L, and jR1R3 Monomers and Dimers. A) Schematics illustrating the relationships between the junction spanning peptides used in the simulations and the Tau187 constructs used in the cellular seeding assays. On the right, junction spanning peptides representing the residues 295(264)-313 in the 2N4R numbering scheme are shown. These regions are highlighted with dotted boxes in the Tau187 constructs used in the study (indigo for jR1R3 and purple for jR2R3). For each corresponding Tau187 construct, a red x or a green check mark annotates if jR2R3 P301L was able to induce puncta in the respective cell line. For 3R constructs the P270L mutation corresponds to the P301L mutation seen in 4R isoforms. B) Illustration of two most populated clusters from REMD simulations. Cluster 1 (Top, Black) highlights hydrogen bonds between aa300-aa305 and K298-Q307. Cluster 2 (Bottom, Cyan) highlights hydrogen bonds between aa300-aa307 and K298-V309. For residues with different numbering in 3R and 4R isoforms, the alternate numbering is shown in parenthesis. Rose colored balls and black colored balls represent the 4 residues that are used to construct the energy landscapes in (C). C) Energy landscapes of α-carbon distances K298-Q307 and aa300-aa305 for (from left to right) jR1R3, jR2R3 (jR2R3 = jR1R3 4Mut), jR1R3 3Mut, and jR2R3 P301L. The y-axis represents the distance between aa300-aa305 (rose), while the x-axis represents the distance between K298-Q307 (black) and are colored as in (B). Clusters 1 and 2 from (B) are annotated in leftmost panel (jR1R3) as dotted lines. Mutations that transition the jR1R3 peptide to the jR2R3 peptide diminish the energy wells stabilizing the hydrogen bonding in Clusters 1 and 2 (Green arrow). This effectively turns on 3R aggregation. The mutation S305K in the jR2R3 construct re-establishes the barriers and turns off 4R aggregation (Red arrow). D) Monitoring the formation of two pairs of hydrogen bonds as jR2R3 transitions into jR1R3 through sequential amino acid swaps. Cluster 1 is represented by black boxes and Cluster 2 is represented with cyan boxes. Cluster 1 with a neutral K305 is shown as an unfilled black box. The bars represent 67% confidence intervals. Protecting (Red) and deprotecting (Green) mutations are highlighted as arrows and represent the mutations shown in panels (C) and (E, F). E) jR2R3 P301L added to cell lines expressing Tau187(3R) P270L did not form aggregates (Left), but when added to cells expressing Tau187(3R) 4mut P270L formed aggregates (Right). F) jR2R3 P301L added to cell lines expressing Tau187(4R) P301L formed aggregates (Left), but when added to cells expressing Tau187(4R) S305K P301L no aggregates were observed (Right).

We turned to replica exchange molecular dynamics simulations (REMD), a widely used method that enhances conformational sampling of proteins by overcoming otherwise inescapable energy barriers and thus exploring extensive free energy landscapes. How might jR1R3 avoid aggregation? A clustering analysis of the replica exchange simulations of jR2R3, jR2R3 P301L, and jR1R3 identified Cluster 1 and Cluster 2 as containing the most prevalent conformations (Figure 3B). In our molecular dynamic simulations, we’ve observed two distinct types of backbone H-bonds in both Cluster 1 and Cluster 2 that help stabilize a β-hairpin conformation, specifically with the GGG motif at the turn. For ease of discussion, we’ve termed the H-bond nearer the GGG turn motif as the “pinch”. This “pinch” appears to keep the hairpin structure closely bound. On the other hand, the H-bond situated closer to the ends of the molecule, which prevents them from unraveling, is termed the “clamp”. Cluster 1 has an H-bond pinch between V/Q300 and S/K305 and an H-bond clamp between K298 and Q307; Cluster 2 has a H-bond pinch between V/Q300 and Q307 and a H-bond clamp between K298 and V309. These H-bonds appear to maintain the hairpin shape in a manner that occludes the amyloidogenic PHF6 sequence.

We quantified the hairpin unfolding free energy landscapes from the replica exchange simulations of jR2R3, jR2R3 P301L, and jR1R3 as a function of the two pairs of diagnostic “pinch” and “clamp” α-carbon distances (Figure 3C). In these landscapes, blue represents low energy, highly populated states, while red represents higher energy, lowly populated states. The most populated conformation (Cluster 1) is in the bottom left of the jR1R3 landscape, while Cluster 2 is slightly above it (both represented with dotted lines) and both have large energy barriers surrounding them. The landscape features distinct minima at short distances indicative of stable conformations. Interestingly, jR1R3 has a significantly larger free energy barrier to unfold than jR2R3 or jR2R3 P301L, and jR1R3 lacks a clear pathway to escape the intramolecularly H-bonded Cluster 1 and Cluster 2. The higher escape energy of jR1R3 may prevent its fibrilization, even in the presence of heparin, by protecting the amyloidogenic PHF6 sequence. This observation is consistent with the requirement for filaments to form, the intramolecular backbone H-bonds of the β-hairpin must be replaced with intermolecular backbone H-bonds as occurs in the disease state (Figure S4A, see orange dashes).

The greater tendency of jR2R3, jR2R3 P301L compared to jR1R3 to form intermolecular H-bond formation, was demonstrated by REMD simulations of dimers. jR1R3 formed significantly fewer intermolecular H-bonds compared to jR2R3 and jR2R3 P301L (Figure S4B). Specifically, conformations with five or more intermolecular H-bonds are rare when either of the monomers exhibits the H-bond pattern of Cluster 1 or Cluster 2. Based on this observation, we conclude that the intramolecularly H-bonded structures of jR1R3, jR2R3 and jR2R3 P301L in Cluster 1 and Cluster 2 conferred resistance to aggregation.

These simulations suggested that an extensive intramolecular H-bond network can safeguard 3R tau containing the junction sequence jR1R3 from aggregation in 4R tauopathies. While similar H-bonds might exist in the jR2R3 region, these are far less stable and have lower energy barriers, facilitating transitions to fibril competent conformations. We performed the *in silico* mutation series of each of the four amino acid differences that converted jR1R3 to jR2R3 in a stepwise manner. Simulating these fragments with explicit-water REMD to observe differences in H-bonding, electrostatic interactions, and hydrophobic interactions revealed that each successive mutation of jR1R3 weakened the protective H-bonding network, as measured by the populations of Cluster 1 and Cluster 2 (Figure 3D). Each of the four residues that differ between jR1R3 and jR2R3 modulated the extent of intramolecular hydrogen bonding and consequently protects PHF6 from aggregation more effectively in jR1R3 than jR2R3. Thus, the gradual weakening of the H-bonds made the peptide more prone to forming intermolecular H-bonds and ultimately aggregation.

The next experiment reversed the direction by mutagenizing jR2R3 to more closely resemble jR1R3 (Figure 3A). Remarkably, in this case, a single mutation in jR2R3 to jR2R3 S305K (a K in jR1R3 occurs in this position rather than an S) drastically increased the free energy barrier to monomer unfolding (Figure 3C, jR2R3 S305K), increased intramolecular H-bonding, and would be predicted to prevent tau aggregation. By simulating the fragments jR2R3 and jR2R3 S305K with REMD, we observed that the S305K mutation approximately doubled the population of the aggregation-resistant Cluster 1 (Figure 3D). K305’s hydrophilic side chain amine tends to orient outward to the solvent, which orients its backbone carbonyl inward where it can form an H-bond with residue 300. We simulated a deprotonated lysine and saw that the doubling of the Cluster 1 population was undone Figure 3D), thus supporting our hypothesis that the charge facilitates the correct orientation of the K305 H-bond.

### Experiments Validate of 4R Energy Barrier Simulations

To validate these computations, a series of 3R tau constructs, with the four sequential mutations within the 1R-3R junction to match those found in the 2R-3R junction of Tau187- P301L, were introduced (Figure 3A Left). In essence, we made a 3R Tau187-P301L that replaced four amino acids of the first repeat with those of the second repeat resulting in a 3R tau that resembled a 4R tau. These constructs were labeled mClover3-Tau187(3R)-P270L to indicate the corresponding site in the 1R-3R sequence with the Tau187-P301L mutation in the second repeat. The mutations were mClover-Tau187(3R)-P270L-E264D, mClover3-Tau187(3R)-P270L-E264D- L266I, mClover3-Tau187(3R)-P270L-E264D-L266I-Q269V, and mClover3-Tau187(3R)-P270L-E264D-L266I-Q269V-K274S. We introduced jR2R3 P301L fibrils to cell lines stably expressing each construct and aggregates were only observed in the construct with all four mutations, i.e. the Tau187(3R) constructs with the jR1R3 segment mutated to mimic the jR2R3 segment (Figure 3E). In other words, a 3R-like tau could be made to aggregate like 4R tau by mutations targeting only the short jR2R3 region. This outcome validates our simulations and suggests that the local structure around the exposed PHF6 motif plays a crucial role isoform-selective seeded aggregation. Thus, jR2R3 P301L induced fibrils rely on a specific set of residues to enable seeding of 4R tau due to exposure of the PHF6 sequence.

While these 4R mutations altered the resistance of 3R tau to aggregate so that it became competent to be recruited to jR2R3 P301L 4R specific tau aggregation, the next question was whether we could prevent 4R aggregation by introducing a mutation into the second repeat that made its structure more similar to 3R tau, and hence resistant to the induction of jR2R3 P301L- induced tau aggregation. In position 305, immediately amino to the VQVII amyloidogenic sequence, a serine is present in the R2-R3 junction, and a lysine is present in R1-R3 junction.

Remarkably, when jR2R3 P301L fibril seed was introduced to a cell line expressing Tau187-P301L-S305K, no significant aggregation was seen (Figure 3F) consistent with the predictions from the REMD simulations. As expected, neither Tau187(3R) nor Tau187(3R)-P270L mutant could be seeded by jR2R3 P301L fibrils in any of our cell culture experiments. This experiment suggests that the single S305K mutation could serve as a prevention of 4R tauopathies.

### Binding of Z70 Nanobody Inhibits Peptide-induced Seeding

We used the previously described camelid heavy-chain-only antibody VHH Z70(18) with a binding sequence that is co-extensive with the PHF6 sequence. VHH Z70 forms five intermolecular hydrogen bonds between VHH Z70 and PHF6. The PHF6 sequence, derived from the structure of disease fibrils, aligns with the crystallized Z70 bound to a free PHF6 peptide (Figure 4A and 4B) (29). Cell lines stably expressing mClover3-Tau187-P301L were transiently transfected with VHH Z70-mCherry and subsequently exposed to jR2R3 P301L fibrils. Twelve hours later, cells were imaged. Analysis in both the mClover-Tau187-P301L and mCherry-VHH Z70 nanobody channels revealed that cells expressing the camelid nanobody contained significantly fewer aggregates, compared to cells lacking mCherry-VHH Z70 (11 ± 7% vs 34 ± 10% of cells, *p* = 2.3 x 10^-8^, Figure 4C, 4D). These results suggest that VHH Z70 can inhibit jR2R3 P301L-seeded aggregation of Tau187-P301L, similar to the seeding experiment using microtubule binding region tau peptide fibrils in HEK293T biosensor lines (18). This, again, confirms that the jR2R3 P301L fibrils serve as mini-4R-tauopathy specific seeds by directly interacting with the PHF6 region.

**Figure 4:**
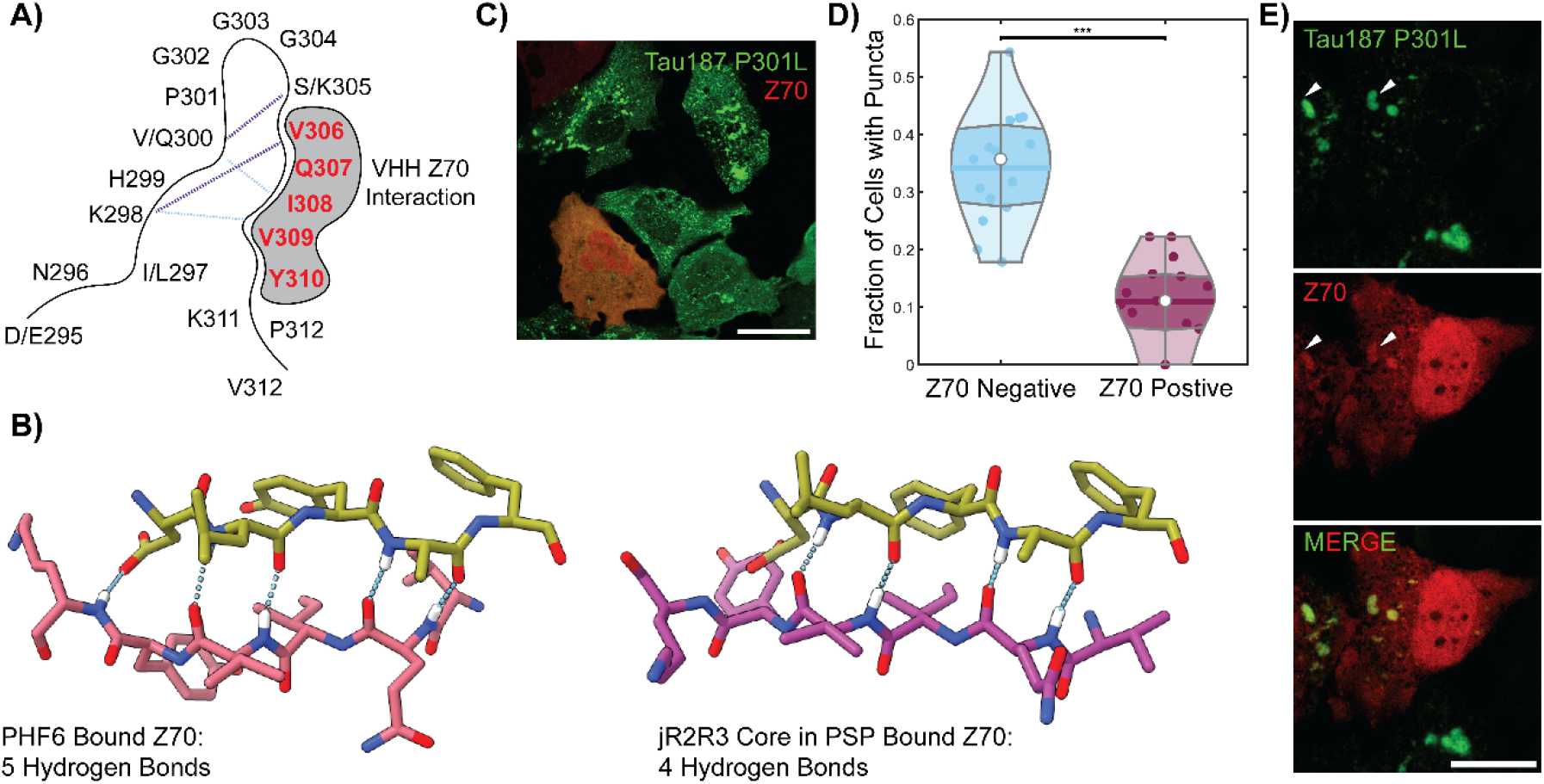
Investigation the Effect of VHH Z70 Nanobody on jR2R3 P301L-Induced Tau Fibrilization in Cells. A) Schematic representation of the SLS hairpin and its most populated hydrogen bonds as determined with molecular dynamics (Cluster 1 in purple and Cluster 2 in cyan dashed lines). The interaction with ZHH Z70 is highlighted by gray shading and its interaction site, the PHF6 motif is represented in red. B) The crystal structure (PDB:7QCQ(17)) of VHH Z70 in olive bound to PHF6 in rose reveals five hydrogen-bond contacts between the molecules (Left). An alignment of the PHF6 sequence from the PSP fibril (PDB:7P65(32), purple, Right) with that bound in the crystal structure of VHH Z70 shows an RMSD 0.848, with four potential hydrogen bond contacts, suggesting that the VHH Z70 nanobody could recognize 4R tauopathy fibrils. C) jR2R3 P301L sonicated fibrils were introduced to stable H4 cell lines expressing mClover3-Tau187-P301L, with a subset of cells transiently expressing a nanobody-mCherry fusion against PHF6 (VHH Z70) in red. Notably, cells expressing VHH Z70 generated fibrils less frequently than those that did not express VHH Z70. Scale bar: 30 μm. D) Quantification of the data from (C) indicates that cells with VHH Z70 formed puncta 11 ± 7% of the time, whereas those without VHH Z70 exhibited puncta in 34 ± 10% of cells, suggesting an inhibitory effect of VHH Z70 on fibril formation. E) Upon addition to preformed aggregates, the VHH Z70 nanobody exhibited clear colocalization with a subset of the tau aggregates, as indicated by white arrow heads. Scale bar: 20 μm.

We next pre-formed cellular aggregates by exposing cells to jR2R3 P301L fibrils first and subsequently transfected cells with mCherry-VHH Z70 in an attempt to clear established aggregates. Under these conditions there was considerable cell death and therefore it was not possible to determine whether aggregates were cleared. However, there were sufficient cells to clearly observe colocalization of the nanobody signal with aggregates (Figure 4E) and demonstrated that the nanobody recognized the aggregated tau populations.

### jR2R3 P301L-Seeding Induces Global Structural Modifications

The variety of cellular studies presented so far showed that the jR2R3 P301L fibrils serve as mini-4R-tauopathey prions that effectively seed naïve Tau187, 0N4R tau, and 2N4R tau constructs that are 10-20 times the length of the jR2R3 P301L fragment. These observations establish that indeed the jR2R3 region is the critical contact point for templated 4R aggregation. The question is then whether a large portion of the tau protein is ordered and incorporated into fibrils when recruited by the jR2R3 P301L seed. To answer this question, we sought to measure the distance changes across tau in the region outside the jR2R3 segment. For this, the seeded material was analyzed by double electron-electron resonance (DEER) experiments to probe the intramolecular fold of tau within the fibrillar aggregates. The distances selected for DEER measurements in Figure 5A were intentionally chosen to showcase regions of Tau187 that are far removed from the immediate vicinity of the jR2R3 P301L peptide and demonstrate the long-range conformational influence exerted by the small peptide, emphasizing its far-reaching templating effects on tau’s structure. A pair of spins were attached to tau at two cysteine residues, with one pair spanning sites 334-360 and another across sites 351-373. The distribution of distances measured between these probes reflects the range of ensemble conformations sampled by the protein. To avoid measuring intermolecular distances, doubly spin labeled tau is mixed with WT tau at a 1:10 ratio. The results then represent a probability density of intramolecular distances across the select pair of spin labels. The specific site pairs that were selected to inform us about the tertiary fold adopted by the seeded tau constructs, e.g., to differentiate between AD, CBD, PSP, and GPT fibrils.

**Figure 5:**
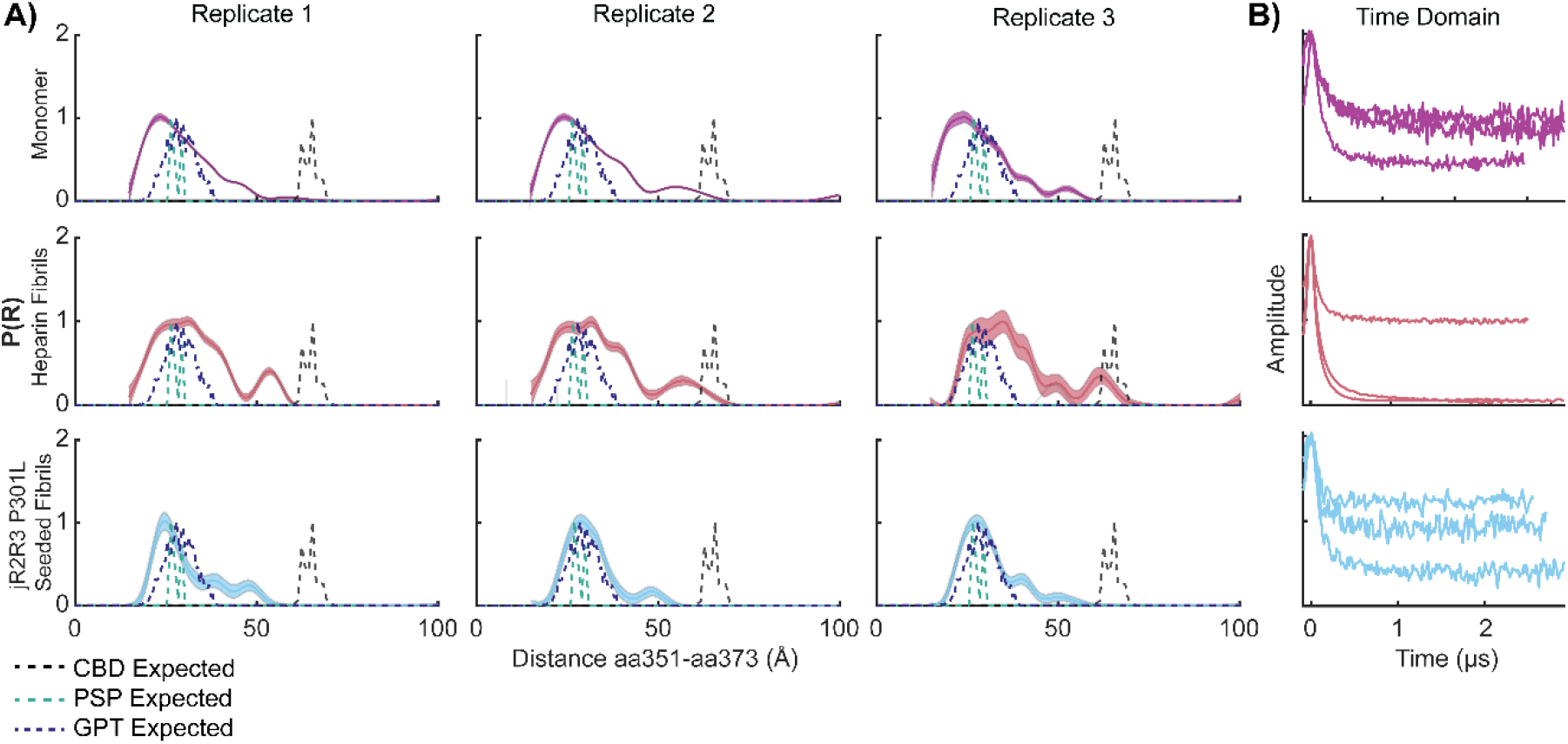
Comparative Analysis of DEER Measurements for Monomeric and Fibrillar Tau Configurations. A) DEER distance distribution plots (P(R)) for the 351-373 spin-pair distances in various tau configurations. The monomer is represented in purple, jR2R3 P301L seeded fibril in cyan, heparin fibril in rose, while the predicted distances for CBD are in black, PSP are in teal, and GPT are in indigo. Triplicates of each condition are plotted across the columns. B) The time domain data for each of the replicates plotted in (A).

The structures formed from jR2R3 P301L fiber-seeded Tau187(4R) material differed substantially from heparin-seeded fibrils (Figure 5A, 5B). Specifically, pairwise distribution plots showed broad distributions for heparin fibrils for both the 334-360 distance (Figure S5A) and the 351-373 distance (Figure 5A). This indicates a sampling of many conformations of fibrils as predicted from the cryo-EM structures of heparin-seeded tau (32). In contrast, jR2R3 P301L- seeded Tau187 fibrils, displayed a narrower distance distribution, indicating a smaller sampled space compared to heparin-seeded fibrils. Interestingly, the 351-373 distance pair, which is predicted to sample widely different distances in PSP, GPT, and CBD fibrils, demonstrated a substantial compaction of the P(R) curve centered at 2.5 nm (Figure 5A). This corresponds to the distance between these two residues in the GPT and PSP cryo-EM structures (29). Unlike the broader peaks observed for both Tau187 monomer and heparin-induced Tau187 fibrils, the peak resulting from the jR2R3 P301L seeded sample is notably narrower and consists of a single, distinct peak, rather than a composite of multiple peaks. Biological triplicate measurements confirmed this observation. Unlike the 351-373 probe pair, the 334-360 probe pair yielded multiple distance peaks that did not correspond closely to either PSP, GPT or the CBD distances (Figure S5A, S5B, S5D, S5E). Instead, they revealed a heterogeneous distance probability distribution that is likely indicative of greater disorder across this distance. Nevertheless, the peaks sampled differed from the heparin-induced distance pair and were narrower and distinct from monomer data. This narrowing of the peaks induced by jR2R3 P301L reinforces the idea that the seed guides Tau187 to a less heterogenous set of structures. Taken together, these data, aligned with cryo-EM structure of jR2R3 P301L fibrils that most closely resembles the GPT distances, suggest that this peptide is capable of templating discrete large structural changes throughout the tau protein and partially captures the GPT distance, a hybrid structure between that in CBD and PSP.

### jR2R3 P301L-Seeded Aggregates have Cross β-strand Structure and an Associated Lipid Signal by Optical Photothermal Infrared Microscopy

Optical photothermal infrared (O-PTIR) microscopy, which provides sub-micrometer resolution infra-red imaging (38–41) was used to probe the structure of jR2R3 P301L-seeded cellular aggregates. In O-PTIR, each pixel of the microscopic image contains an IR spectrum over a defined range of wavenumbers allowing for the investigation of the chemical and structural composition at specific fluorescent structures (Figure 6A, 6B). Measurements of the cellular aggregates revealed a significantly higher cross β-strand content compared to the cytoplasm, as indicated by the substantial difference in the mean values of the ratio of peaks at 1634 cm^-1^ and 1654 cm^-1^ (p = 1.6 x 10^-5^, Figure 6D). This result is consistent with the expected properties of aggregates forming fibril-like structures. Interestingly, a local increase in signal at 1750 cm^-1^ associated with the aggregate was observed, indicative of a lipid signal. When compared to the cytoplasm, the difference in the ratios of 1750 cm^-1^ and 1654 cm^-1^ was also highly significant (p = 7.8 x 10^-7^, Figure 6D).

**Figure 6:**
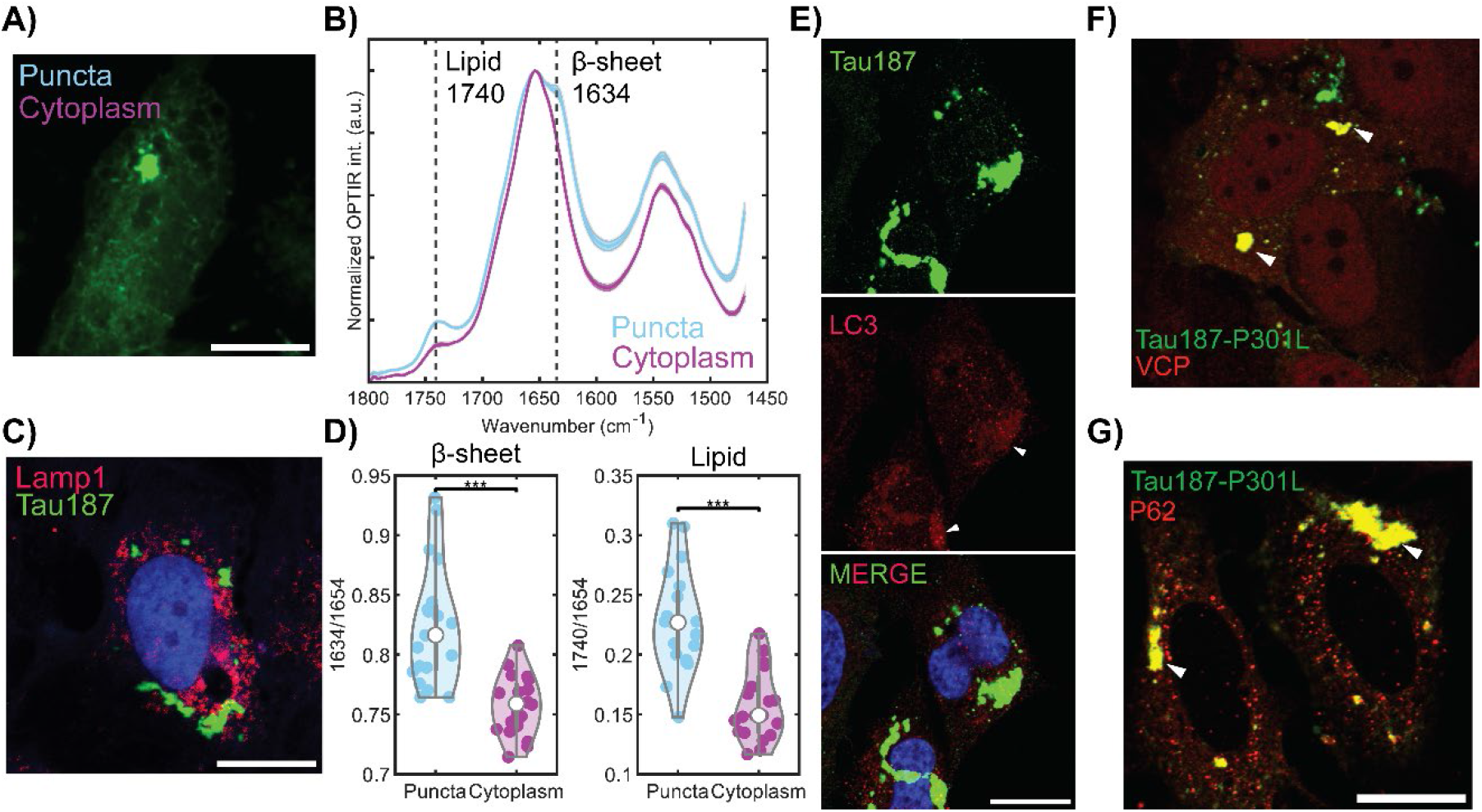
Presence of Lipid in jR2R3 P301L-Seeded Cellular Aggregates using Optical Photothermal Infrared Microscopy and Analysis of Autophagosome and Lysosomal Markers. A) O-PTIR interrogation of cells expressing mClover3-Tau187-P301L. Measurements (depicted as cyan and purple dots) were taken on tau aggregates and cytoplasm, respectively. Scale bar: 15 μm. B) Averaged spectra of measurements from (A), with distinct peaks at 1740 cm^-1^ (lipid) and 1634 cm^-1^ (β-sheet) in aggregates. Error bars represent SEM. C) Representative cell stained with Lamp1 antibody (red) to visualize lysosomes. Tau187 puncta signal (green) shows minimal colocalize with Lamp1 signal. Scale bar: 20 μm. D) Quantification of lipid and β– sheet content from 20 independent measurements in aggregates (cyan) and cytoplasm (purple), for both lipid and B-sheet in aggregates revealing highly significant differences. E) Tau (green) co-localization with LC3 (red) indicates significant overlap with the largest tau aggregates (white arrows), suggesting a possible early stage of autophagy recruitment. Scale bar: 20 μm. E, F) Colocalization of protein degradation markers VCP (red, E) and P62 (red, F) with large tau aggregates (green). Scale bar: 20 μm.

### Cellular Effects of Seeded Tau Aggregates

To examine the possibility that tau aggregates were associated with lysosomes or autophagosomes, cells were stained with either a Lamp1 or a LC3 antibody. Small, punctate signals typical of lysosomes or autophagosome did not substantially co-localized with the aggregate signals (Figure 6C). The Manders’ coefficient for tau to Lamp1 overlap was 0.12 ± 0.1, and for Lamp1 overlap to tau was 0.055 ± 0.05 (N=15 cells), indicating minimal overlap. For tau to LC3, the overlap was 0.41 ± 0.2, and LC3 to tau overlap was 0.44 ± 0.1 (N=8 cells), suggesting modest colocalization. Interestingly, a substantial overlap was observed between LC3 and the largest tau aggregates, with a Manders’ coefficient of 0.90 ± 0.2 for tau overlap to LC3 under specific thresholds that capture only the largest tau aggregates (Figure 6E). Thus, there is some attraction of LC3 to the larger aggregates.

The large perinuclear aggregates of tau protein observed in our jR2R3 P301L-seeded cells hinted at the involvement of cellular degradation mechanisms beyond the capacity of traditional autophagy or proteasomal pathways to clear proteins (42), and thereby necessitate the deployment of alternative clearance strategies such as aggresomes (43, 44). To examine this possibility, we stained cells expressing aggregate induced by jR2R3 P301L with antibodies targeting SQSTM1 (also known as P62) and VCP (also known as P97). Both of these proteins have been implicated in previous studies as colocalizing with tau inclusions, suggesting a potential role in the degradation of tau aggregates (45–52). Immunofluorescence analysis (Figure 6F, 6G) revealed that both SQSTM1 and VCP exhibited pronounced colocalization with the large tau aggregates in the jR2R3 P301L-seeded cells. This observation aligns with existing literature that demonstrates the colocalization of SQSTM1 and VCP with tau inclusions (48–53), suggesting their involvement in the degradation of tau aggregates.

### Dynamics and Heterogeneity of Tau Aggregates

Before the introduction of seeds, all cell lines—including WT and P301L mutants— displayed typical microtubule staining patterns (Figure 2A Left, S2B, S2C). In cells with mature aggregates, which can range in size from very large (up to 20 μm) to minute, diffraction-limited specks (<250 nm), the population of tau bound to microtubules was greatly diminished (Figure 2C). Aggregates larger than 1 μm in size displayed a diverse range of morphologies, with some appearing elongated and others lumped together. This irregularity is reflected in a variable mean circularity value centered at 0.55 ± 0.25. The population of cells with the most aggregates were entirely devoid of tau microtubule staining (Figure 2E) and by four days in culture such cells dominated. In this experiment, cells actively expressing new monomeric tau preferentially partitions into aggregates rather than binding to existing microtubules suggesting that the aggregate has a greater affinity for tau than the microtubules. However, after staining with an anti-α-tubulin antibody, an intact microtubule network remained presumably associated with other microtubule-associated proteins (Figure 7A).

**Figure 7:**
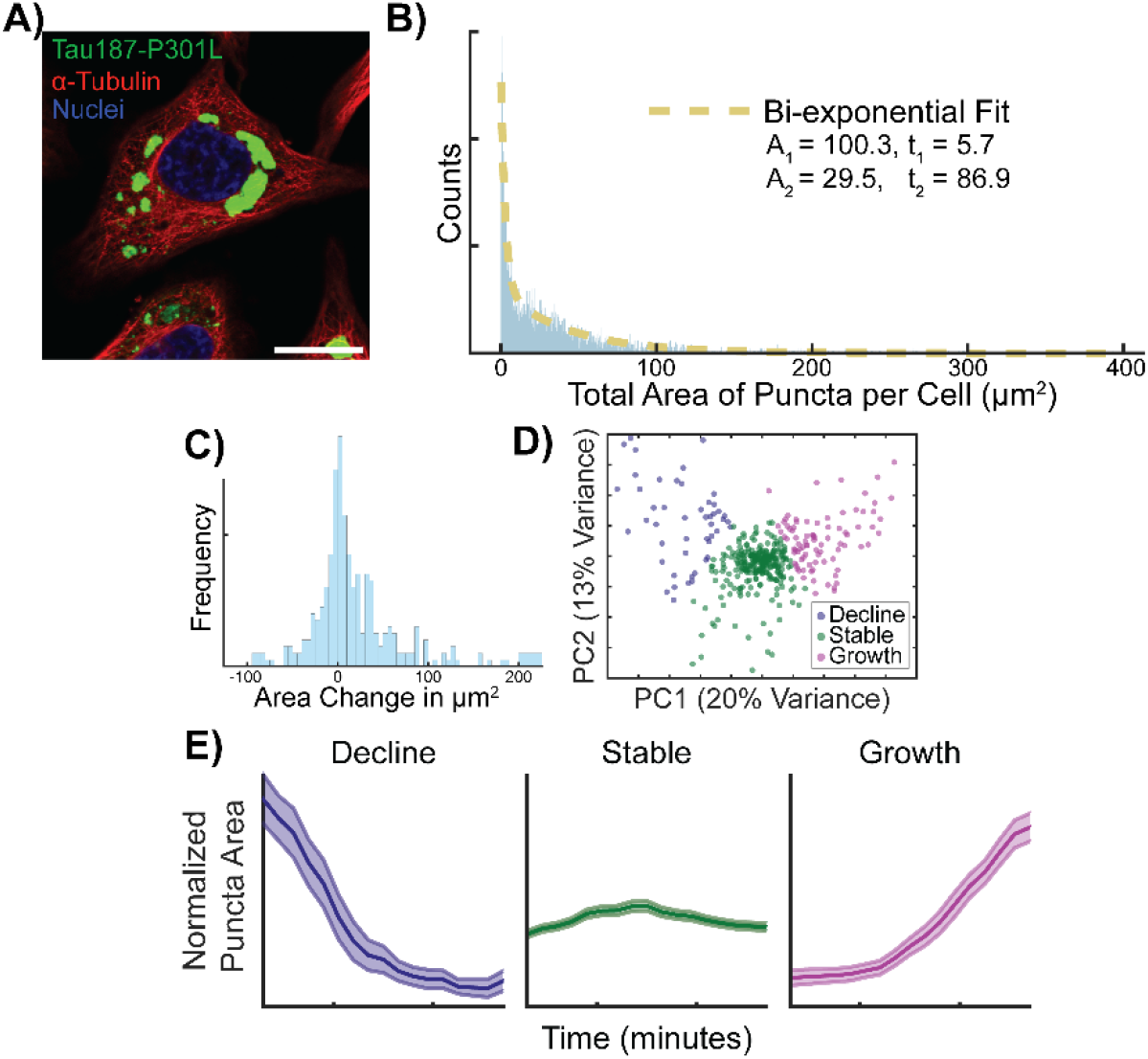
Growth, Stability, and Shrinkage in Puncta. A) Microscopy images showing intact microtubule network (α-tubulin staining in red) despite no visible tau binding to microtubules after large aggregates form (green). Scale bar: 20 μm. B) Histogram of the total puncta area per cell fit to a bi-exponential decay curve (sand dotted line). C) Monitoring cells with jR2R3 P301L-induced aggregates for 10 hours, imaging every 30 minutes. The total change in aggregate area per cell was plotted as a histogram. D) PCA analysis on the trajectories of the changing aggregate area; K-means clustering identified three clusters that represent cells with declining (indigo), stable (green), and growing (purple) aggregate areas. E) The average of the individual trajectories based on K-means clustering are plotted, showing decline, stability, or growth. Shaded area represents the SEM of the normalized, combined trajectories. E) Colocalization of protein degradation markers VCP and P62 (red) with large tau aggregates (green). Scale bar: 20 μm.

To systematically investigate the temporal evolution of tau aggregation, live cells were monitored for a period of 10 hours, with images of tau aggregates captured at 30-minute intervals. Aggregates did not appear to fuse but sometimes clumped together. Fragmentation of the aggregates was not observed. A few aggregates were spherical, but most were irregular in appearance. A few had a linear appearance that resembled the seeded aggregates previously observed in proximity to microtubules (54). Individual cells were tracked, allowing for the quantification of the total area occupied by aggregates, the average size of aggregates, and the number of aggregates per cell. To enhance data reliability, we excluded cells undergoing division or exhibiting signs of cell death.

Our analysis revealed that both the total area occupied by aggregates per cell and the number of aggregates per cell followed a bi-exponential distribution (Figure 7B). We then classified changes in total aggregate area within individual cells and found that the total aggregate area decreased in 48 cells, remained stable in 267 cells, and increased in 86 cells over 10 hours (Figure 7C). We applied K-means clustering analysis to the trajectories of individual cell total area, effectively segregating the data into three distinct subpopulations based on aggregate dynamics (Figure 7D). We plotted these subpopulations and observed clear patterns of decline, stability, and growth in aggregate areas for the respective clusters (Figure 7E).

## Discussion

PHF6 and PHF6* oligomers, rich in β-strand content (12), can form self-complementary beta-sheet pairs known as steric zippers (55), but they are unable to template tau misfolding in cells. By extending the PHF6(*) core to include portions of the tau repeat 2 and 3 sequences, seeding active fibrils can be generated (17, 21, 56, 57). However, a higher order structural motif is needed to orient and hold the PHF6 segment with its hydrogen bond forming backbone moieties oriented along the fibril axis and to achieve shape propagation of a nucleating tauopathy fold. In this study, we ask whether jR2R3 P301L fibrils are active tau prions by showing recruitment of full-length intracellular tau over multiple passages and by achieving isoform-selective seeding. By introducing jR2R3 P301L fibrils into cells, we confirm the fibril, not the monomer, as the tau peptide that can template misfolding. Furthermore, the misfolding and induced conformational changes are seen in regions of tau well outside the 19-residue long jR2R3 segment, supporting our hypothesis that misfolding of a critical motif seeds the aggregation of many naïve tau molecules and induces misfolding to the rest of the tau sequence. The so treated intracellular tau forms sarkosyl insoluble material with properties that parallel human brain tauopathies, including MC1 antibody staining, fibril-like morphologies seen by TEM, and molecular weights that approximate these complex endogenous seed-competent assemblies.

Critical to the design of the jR2R3 P301L peptide is its correspondence to the R2R3 junction sequence, uniquely present in 4R tauopathy fibrils in AGD, PSP, CBD, ARTAG, GGT and GPT (25), resulting in fibrils that require the precise 4R tau sequence to seed intracellular 4R tau. A 4R tau in which an amino acid substitution mimicked a 3R tau sequence could not be seeded. This shows that the location of templating that can differentiate between a 4R tau versus a 3R tau substrate must encompass the jR2R3 junction of the peptide we studied here. To observe such exquisite isoform selectivity between different tau constructs differing by a single residue at the jR2R3 junction, the orientation of hydrogen bond-forming backbone along the fibril axis is an insufficient condition. Rather, there must be a mechanism that aligns the tau monomer substrate on the surface of the tau fibril observed in Vigers et. al, in which a hydrophobic hotspot, hyper-localized around sites 300 and 301, pins the jR2R3 P301L peptide to the fibril end surface, in register.

The degree of ordering and selection extends beyond the region of tau corresponding to the peptide. The probability distribution of intra-molecular distance, P(r), measured by DEER of jR2R3 P301L fibrils show that at least two distinct structures form that are outside the core region identified by cryo-EM. The selection of only one of the peptide fibrils must be achieved to narrow the heterogeneity of the longer Tau187 fibrils seeded by the short jR2R3 peptide. As expected from the structure of the seed peptide, the DEER distances of seeded Tau187 fibrils were consistent with values expected from PSP or GPT. The selection of induced structures from the vast disordered landscape was more restricted than in heparin-induced fibrils (Figure 5A, S5A), but without the extraordinary fidelity of tau templating in the brain that generates one or very few tau structures.

The most salient feature of prion-like templating by jR2R3 P301L is its specificity for 4R tau. Such specificity must be rooted in a distinctive shape of the active fibril end that serves as a molecularly precise template and is thermodynamically capable of recruiting naïve tau, with its PHF6 and its flanking domain in register to achieve templating. By substituting specific amino acids in the flanking sequence that will weaken or strengthen access to the PHF6 region, 4R tau can acquire or lose competence to initiate a 4R tauopathy fold. These substitutions are derived from those amino acids in jR1R3 that depart from jR2R3 and were inferred from the REMD. The simulations showed clear energetically accessible pathways from a shielded loop region due to H-bonding to a more accessible internal loop structure in jR2R3 that is prone to intermolecular H-bonds and hence oligomerization. The single amino acid substitution, S305K, that can prevent jR2R3-induced aggregation, suggested a site where a single codon substitution could prevent 4R tauopathies. As a point of background information, S305 is the last codon of MAPT exon 10, and is included in the RNA hairpin loop that regulates exon 10 alternative splicing. Although mutations have been identified in this codon that increase 4R tau, like all other splice site mutations, none of mutations correspond to S305K used here for its effects on the tau protein (58).

Three factors potentially explain why the S305K mutation inhibits 4R tau aggregation: steric hindrance of the known 4R tau folds, increased hydration at the dewetting interface, and altered intramolecular H-bonding within monomers. Among four solved 4R tau folds, three—CBD, PSP, and GGT—have an S305 oriented inwardly within the strand-loop-strand motif (Figure S6). A bulky, inwardly-pointing K305 side chain would sterically disrupt the side chain zipper, preventing these structures from forming. In contrast, a fourth 4R tau fold, termed “GPT”, presents an S305 that faces outward from the strand-loop-strand motif, leaving this motif undisturbed by the bulky K305. However, even in this context, a K305 side chain would sterically disrupt contacts with I354 and L357 potentially impeding fibrillization. Beside steric considerations, the S305K mutation might disrupt aggregation due to its positively charged side chain which interacts favorably with water. Consequently, this side chain and the adjacent residues will be more resistant to dewetting, which is an essential step in nucleation and elongation. Recently, time-resolved cryo-electron microscopy detected a “first intermediate” on-pathway to Alzheimer and CTE fibrils consisting of an ordered core spanning residues 302-316 (59). Interestingly, the AD/CTE-precursor region includes the PHF6 region and C-terminal extension, but not the complete counter strand needed to form the SLS core of 4R tauopathy fibrils. This small shift relative to jR2R3 (295–313) may be enough for the region 302-316 to follow a different folding pathway and conformational rearrangement to ultimately generate fibrils resembling the 3R/4R tauopathy forms as observed in AD filaments.

Although the peptide here was derived from cryo-EM structures of tau, its relationship to *in vivo* proteoforms is unknown. Many small tau fragments have been reported in tauopathies; however, their role in disease is debated. The 19-amino-acid sequence corresponds to a relatively “cleavage poor” region of tau (60) possibly related to the two histidines which is the only amino acid with a pKa within the physiological range and will make the peptide uniquely suited to be protonated inside the acidic lysosome. While mainly of interest as a heuristic probe, the resistance of the peptide to proteolysis under physiologic conditions may generate a similar peptide *in vivo*.

Many intracellular factors can lead to the nucleation and propagation of such tau peptides, as long as they contain amyloidogenic domains. The exposure to and compaction of the cellular space by lipid droplets or in some other entity may force the formation of hydrophobic bonds with an aqueous solvent and intermolecular hydrogen bonds found in tau aggregates. Solvent-exposed hydrophobic regions can favor interactions with low-molecular-weight hydrophobic or amphiphilic lipids known to interact with tau and induce aggregation (61). Within intrinsically disordered proteins, lipid-binding occurs in more structured regions, such as the relatively more ordered repeat region in tau where its prion properties reside (62). Indeed, using solid-state NMR and transmission electron microscopy, a tau construct encoding the microtubule-binding repeats and a proline-rich domain was reconstituted into cholesterol-containing phospholipid membranes in a transition from a random coil to a β-sheet conformation over weeks (63). Cellular aggregates displayed colocalization with VCP and SQSTM1, proteins essential for aggresome formation and selective autophagy (47–49, 53, 64). SQSTM1, known to interact with tau and autophagosomes, has diminished expression in Alzheimer’s disease (43) and its depletion leads to hyperphosphorylated tau accumulation (50). Meanwhile, VCP aids in the degradation of tau aggregates (49) and is crucial for proper aggresome formation (65), implicating a critical role for both proteins in the pathogenesis of tauopathies.

These studies suggest several entry points for 4R tauopathy interventions involving either nanobodies or small molecules, but all pointing to the junction spanning U-shape R2R3 structure with its relative exposure of the PHF6 amyloidogenic sequence capable of templating the fibril end surface. Structural studies suggest a similar rationale might be applied to the amyloid-beta-peptide (66). Furthermore, the dynamic size changes of the aggregates observed intracellularly may offer some potential for disaggregation approaches.

## Materials and Methods

### Preparation of Purified Tau Constructs

Peptides obtained from Genescript, all featuring uncapped termini, were dissolved in ddH_2_O and stored immediately as 100 μM aliquots at -80°C. The aggregation process was facilitated in a 50 mL Eppendorf tube where the stock peptide was reduced to a 50 μM concentration using 20 mM HEPES, pH 7.4, accompanied by a 4:1 ratio addition of 200 μM heparin (average molecular weight 16 kDa). Continuous shaking at 200 rpm in a 37°C incubator was performed for 24 hours. The resultant fibrils were isolated using a 50 kDa cutoff concentrator and rinsed 3 times with MilliQ water to remove excess peptide and monomeric peptide. The fibrils were then lyophilized overnight using a FreeZone 2.5 Liter -84C Benchtop Freeze Dryer. Upon quantification of the fibril mass, we reconstituted the seed stock to 1 mM using 20 mM HEPES, pH 7.4, and froze it at −80°C.

Tau187 expression and purification procedures followed previously reported methods^67^. We transformed E. coli BL21 (DE3) cells with various plasmids and stored the cells as frozen glycerol stocks at −80°C. An overnight growth of cells in 10 mL Luria Broth preceded the inoculation of 1 L of fresh LB. The incubated the cells at 37°C at 200 rpm, supplementing with 10 μg/mL kanamycin, until an optical density at λ = 600nm of 0.6–0.8 was achieved. We induced expression by introducing 1 mM isopropyl-ß-D-thiogalactoside for 2–3 h. Post expression, cells were harvested via centrifugation at 4500 g for 20 min.

Harvested cells were resuspended in lysis buffer (Tris-HCl, pH = 7.4, 100 mM NaCl, 0.5 mM DTT, 0.1 mM EDTA) supplemented with 1 Pierce protease inhibitor tablet per 5 mL lysis buffer, 1 mM PMSF, 2 mg/mL lysozyme, 20 μg/mL DNase and 10 mM MgCl2, and incubated on ice for 30 min. The samples underwent three freeze-thaw cycles using liquid nitrogen, followed by 10-minute centrifugation at 10,000 rpm to discard cell debris. Another, 1 mM PMSF was added, and samples were then heated at 65°C for 12 minutes and cooled on ice for 20 minutes before undergoing another round of centrifugation to remove the precipitant.

The clear supernatant was incubated with pre-equilibrated Ni-NTA resins overnight in buffer A (20 mM sodium phosphate, pH = 7.0, 500 mM NaCl, 10 mM imidazole, 100 μM EDTA). We subsequently transferred the resins to a column and washed with 20 mL of buffer A, 25 mL buffer B (20 mM sodium phosphate, pH = 7.0, 1 M NaCl, 20 mM imidazole, 0.5 mM DTT, 100 μM EDTA). We collected the purified protein through elution with 15 mL of buffer C (20 mM sodium phosphate, pH = 7.0, 0.5 mM DTT, 100 mM NaCl, 300 mM imidazole). SDS-PAGE confirmed the purity of the fractions. After buffer exchange into a DTT-free working buffer (20 mM ammonium acetate, pH 7.0), we immediately froze the proteins and stored them at −80°C.

### Cell Seeding Experiments

H4 neuroglioma cells stably expressing either mClover3-Tau187-WT, mClover3-Tau187-P301L, mClover3-0N4R-WT, mClover3-0N4R-P301L, mClover3-2N4R-WT, or mClover3-2N4R-P301L were cultured in DMEM, supplemented with 10% FBS, 100 μg/ml penicillin/streptomycin, and 500 μg/ml geneticin. The cell cultures were maintained in a humidified atmosphere of 5% CO_2_ at 37 °C. For seeding experiments, cells were plated in 96-well plates at a density of 20,000 cells per well. On the following day, cells were transfected with various tau species (at a final concentration of 2 μM unless stated otherwise) using 1.25 μL of Lipofectamine 2000 (Thermo Fisher) per well. The tau seeds were subjected to sonication for 30 seconds using the microtip of a Qsonica sonicator, operating at a 30% duty cycle. For the quantification of the assay, cells containing one or more puncta were counted and normalized by dividing by the total number of cells in each well. Each well in the cultured plate contributed one data point, and the quantification results were obtained from at least five independent cultures.

For experiments assessing the relative incorporation of different tau isoforms, H4 cell stably expressing mRuby3-Tau187-P301L-P2A-mClover3-Tau187-P301L, mRuby3-Tau187(3R)-P2A- mClover3-Tau187-P301L, mRuby3-Tau187(3R)-P2A-mClover3-Tau187(3R), were cultured following the aforementioned protocol.

### Fluorescence-guided OPTIR imaging setup, spectral measurement, and data processing

OPTIR spectroscopy and co-registered fluorescence imaging were performed on a mIRage LS microscope (Photothermal Spectroscopy Corp.). The mid-IR laser used as the photothermal pump source was pulsing at 100 kHz repetition rate with 1% duty cycle. The visible probe source was a continuous wave laser with a center wavelength of 532 nm. The counter-propagation geometry of the mid-IR and the visible beam was used. The mid-IR beam was focused on the sample plane below the sample substrate (CaF_2_) with a reflective objective (40x, 0.78NA, Pike Technologies), and the visible beam was focused on the sample plane from the top with a refractive objective (50x, 0.8NA, Olympus). Epi-detected light was collected and focused on a photodiode. The OPTIR signal was demodulated with a lock-in amplifier at 100 KHz. The system was enclosed and purged under gentle nitrogen flow to minimize water vapor interference of the spectra interpretation. For co-registered widefield fluorescence imaging, a filter cube set suitable for mClover3 excitation and emission was used. Typical power at the sample for mid-IR and visible were a few mW depending on the sample locations.

To prepare for OPTIR spectroscopy measurements, the cells were rinsed twice with deionized water before drying in the air. For the data acquisition, we first acquired widefield fluorescence imaging to reveal the tau distribution in cells. We then pinned to puncta regions and puncta-free regions in cells to acquire OPTIR spectra. The spectra spanned 1480 to 1800 cm^-1^ (5.55 to 6.75 μm), covering protein amide II, protein amide I, and lipids signal. All raw spectra were normalized with mid-IR power spectra to reveal the sample-related IR peaks. For the OPTIR spectra presented in the manuscript, OPTIR spectra were normalized to OPTIR intensity at 1654 cm^-1^. For secondary structure quantification, β-sheet to α-helix ratio was calculated using OPTIR intensity at 1634 cm^-1^ and 1654 cm^-1^. Lipid contents were quantified by normalizing OPTIR intensity at 1740 cm^-1^ to 1654 cm^-1^. For each cell, we acquired at least 8 spectra at different puncta and puncta-free locations, where the mean intensity of the ratioed value was used to represent a single cell. Ten cells were used to perform statistical analysis.

## Acknowledgments

We acknowledge support by the NIH grant 5R01AG056058-07 and the Rainwater Foundation. The authors acknowledge support from the Center for Scientific Computing at the California Nanosystems Institute (CNSI, NSF grant CNS-1725797) for the availability of high-performance computing resources and support. This work used the Extreme Science and Engineering Discovery Environment, which is supported by the National Science Foundation grant ACI- 1548562 (MCA05S027). J.E.S acknowledges support from the NSF (MCB-1716956). We acknowledge the use of the NRI-MCDB Microscopy Facility and the Resonant Scanning Confocal supported by NSF MRI grant DBI-1625770.

