## Supplementary Methods and Figures for "Precision Proteoform Design for 4R Tau Isoform Selective Templated Aggregation"

**This PDF file includes:**

Extended Methods
Figures S1 to S6
SI References

**Other supporting materials for this manuscript include the following:**

**Extended Methods**

***Cloning of Tau Constructs***

We assembled mClover3-2N4R-WT constructs using the HiFi Reaction Kit from NEB. Additionally, we generated various isoforms such as mClover3-0N3R-WT, mClover3-0N4R-WT along with truncations including mClover3-Tau187(4R)-WT, mClover3-Tau187(3R)-WT through deletion PCRs and KLD reactions using NEB kits.

Using site-directed mutagenesis, we created mutant constructs, specifically mClover3-Tau187-P301L, mClover3-0N4R-P301L, and mClover3-2N4R-P301L. Additionally, using similar techniques, we created mClover3-Tau187(3R)-P270L, mClover3-Tau187(3R)-P270L-E264D, mClover3-Tau187(3R)-P270L-E264D-L266I, mClover3-Tau187(3R)-P270L-E264D-L266I-V269Q, and mClover3-Tau187(3R)-P270L-E264D-L266I-V269Q-S274K

Constructs with dual tau expression, such as mRuby3-Tau187-P301L-P2A-mClover3-Tau187-P301L, mRuby3-Tau187(3R)-P2A-mClover3-Tau187-P301L, and mRuby3-Tau187(3R)-P2A-mClover3-Tau187(3R), were synthesized using HiFi reactions with the previously mentioned constructs as templates.

We procured pET28a-Tau187-WT-His and pEt28a-Tau187-P301L-His constructs from Genescript and constructed pET28a-Tau187(3R) using deletion PCR.

For DEER experiments, we generated Tau187 Cysless constructs with one or two cysteine mutations using site-directed mutagenesis.

***Thioflavin T Experiments***

Thioflavin (ThT) experiments were conducted using a BioTek Synergy 2 fluorescent plate reader. Each well of a 384-well plate (Corning, low volume non-binding surface, black with clear flat bottom) contained 50 μM of SLS or Tay monomer, 20 μM ThT, 12.5 μM heparin or 12% seed 20 mM HEPES HEPES buffer (pH 7.4), making a total volume of 20 μL per well.

The plate reader was set to a temperature of 37°C and allowed time to equilibrate. Subsequently, ThT fluorescence intensity was recorded at excitation and emission wavelengths of 440 nm and 485 nm, respectively. Measurements were taken every 2 minutes until a plateau in fluorescence intensity was observed. Each experiment was performed with five replicates and repeated three times using independent samples, typically spanning a total time of 48 hours.

***TEM Analysis***

For transmission electron microscopy (TEM) analysis, 5 μL of recombinant tau fibril samples were placed onto a glow-discharged copper grid (Electron Microscopy Science, FCF-200-Cu) for 20 seconds before blotting dry with filter paper. The samples were then stained by applying 5 μL of 1.5% (w/v) uranyl acetate solution, immediately blotting dry. An additional 5 μL of uranyl acetate solution was added for 60 seconds, followed by blotting dry. The samples were examined using a Thermo Scientific Talos G2 200X TEM/STEM microscope operating at 200 kV and room temperature. The grids were imaged using a Ceta II CMOS 4k x 4k camera at the specified magnifications.

***Object Segmentation and Tracking***

All image processing was carried out using Python 3.10.12. The time series of cell images featuring puncta were loaded utilizing the OpenCV-python 4.7.0.72 package. The analysis of these images was conducted using scikit-image 0.19.3 package.

Initially, nuclei were identified by applying Li thresholding (1) to the Hoechst intensity image. Objects with size smaller than 100 $\mu m^2$ were excluded from further analysis as noise, and the remaining connected regions were labeled as separate nuclei.

Subsequently, the mClover3-Tau187 fluorescence intensity images, after being subjected to blurring, were used for the detection of cell bodies through mean thresholding (2). The masks for nuclei and cell bodies were combined to form complete cell masks. The watershed algorithm was employed, using labeled nuclei as seeds, to delineate individual cells.

Cells obtained through this process were tracked across successive frames using centroid tracking. In this approach, a centroid in a new frame was associated with the nearest centroid in the preceding frame, marking them as the same object.

Puncta were detected using the Laplacian of Gaussian blob detection method. In each frame, puncta located within the mask of a cell object were assigned to that cell.

***Computational Modeling***

*REMD Simulations*

Replica-exchange molecular dynamics (REMD) simulations were carried out using the Gromacs package (versions 2019.6 and 2020.1) (3,4). Simulations were conducted in rhombic dodecahedral boxes of approximately 7.5 nm length, comprising 19-mer peptide(s), around 7860 TIP4P-D water molecules, and neutralizing chlorine ions. Peptides were simulated with zwitterionic termini. In the case of monomer simulations, the peptides began in extended conformations. For dimer simulations, the peptides were initialized in the two most common monomer conformations based on a Daura clustering analysis with a 4Å RMSD cutoff, and they were separated by a couple of nanometers.

The force field AMBER99SB-disp (5) was used, and periodic boundary conditions were applied along all three box vectors. The simulations included an initial 5 ns run in the NPT ensemble at 1 bar pressure and 300 K. The v-rescale thermostat (0.5 ps time constant for temperature coupling) and the Parrinello-Rahman barostat (3 ps time constant for pressure coupling) were used for this step. Next, 5 ns long NVT simulations with the Nose-Hoover thermostat (3 ps time constant for temperature coupling) were performed at 300 K temperature using the average box-size from the final three ns of the previous NPT step (6,7). Throughout the simulations, the Particle Mesh Ewald (PME) method was used for calculating the electrostatic interactions with a grid-spacing of 0.12 nm (8). A time step of 2.5 fs was employed using a leap-frog integrator (9). A cutoff distance of 1.2 nm was set for all nonbonded interactions. For simulations excluding MTSL, the LINCS algorithm was used to constrain bonds with hydrogens (10). Water molecules were maintained fully rigid using the SETTLE algorithm (11).

Subsequently, 60 replica systems were created at temperatures ranging from 300 K and 456 K: The temperatures of the replicas were: 300.0, 301.9, 303.8, 305.7, 307.7, 309.7, 311.7, 313.7, 315.8, 317.9, 320.0, 322.1, 324.3, 326.5, 328.7, 330.9, 333.2, 335.5, 337.8, 340.1, 342.5, 344.9, 347.3, 349.7, 352.2, 354.7, 357.2, 359.7, 362.3, 364.9, 367.5, 370.1, 372.8, 375.5, 378.2, 380.9, 383.7, 386.5, 389.3, 392.1, 395.0, 397.9, 400.8, 403.7, 406.7, 409.7, 412.7, 415.8, 418.9, 422.0, 425.2, 428.4, 431.6, 434.9, 438.2, 441.6, 445.0, 448.5, 452.0, 455.6 K. Each replica underwent a 5 ns NVT equilibration run per without exchanging replicas.

Following this, the replicas were subjected to 5 ns equilibration runs with replica exchanged occurring every 3 ps, achieving an average exchange probability of about 30%. This was followed by production REMD simulations ranging from 500-750 ns per replica. The initial 40 ns were discarded for further equilibration, and certain simulations employed longer equilibration times as discussed in the next paragraph.

Equilibration times were determined using a heuristic that maximizes the number of effectively uncorrelated measurements. The autocorrelation function was integrated for equilibration times in 10 ns increments starting from 50 ns, and the equilibration time with the most effectively uncorrelated measurements was chosen. Dimer simulation equilibration times ranging from 80 to 140 ns were selected by maximizing the number of effectively uncorrelated intermolecular hydrogen bond counts. Autocorrelation functions were measured using pymbar's timeseries module. Confidence intervals were determined with bootstrapping. Residues were assigned secondary structures using the DSSP algorithm in mdtraj. ChimeraX and Pymol were employed for visualizing conformations. Hydrogen bonding analysis was conducted in MDAnalysis, utilizing distance cutoffs of 3.5 Å and angle cutoffs of 140° for determining hydrogen bonds. Energy landscapes were generated by measuring the 2D probability density, taking a logarithm, scaling by Boltzmann's constant and temperature, and applying Gaussian interpolation to smooth the landscape.

*Water Triplet Angle Analysis*

The water triplet angle distribution refers to the distribution of angles formed by the oxygen of a central water and the oxygens of each pair of its neighboring water molecules. This distribution has been demonstrated to reveal patterns in the structing of water in response to different solutes and interfaces (12,13). In this study, the distribution of hydration water was computed to be 4.25 Å around the heavy atoms of each residue's backbone or side chain. The 4.25 Å cutoff corresponds to the second minimum in the radial distribution function (RDF) between the heavy atoms of the backbone or side chain and the oxygen atoms of water. Neighboring water molecules were defined as those having oxygen atoms within 3.4 Å of the central water molecule. The hydrophilic signature was defined as the fraction of hydration waters around each peptide that form angles between 45-

50° with adjacent waters, normalized by the fraction of bulk waters forming angles between 45-50° with adjacent waters. Conformations were analyzed based on data extracted every 10 ps from REMD simulations. Each frame contained statistically independent water triplet angle measurements, which were based on measured autocorrelation times. Confidence intervals of 67% were determined though bootstrapping with over 1000 independent measurements taken with replacement.

***Spin Labeling of Tau***

Tau187, with its native cysteines mutated to serines and the specified residues mutated to cysteines, was expressed and spin-labeled to investigate distances between two sets of target residues (residues 351 and 373, and residues 334-360). The native cysteines at 291 and 322 were mutated to serines. Additionally, a similar tau construct without any cysteines (referred to as tau-cysless) was expressed and purified. The protein was spin-labelled using MTSL((1-Acetoxy-2,2,5,5-tetramethyl-δ-3-pyrroline-3-methyl) Methanethiosulfonate), which was purchased from Toronto Research Chemicals.

Before the labeling process, the samples were treated with a10x molar excess of DTT to reduce any cysteine residues. The DTT was subsequently removed using a PD-10 desalting column. Subsequently, a 10x molar excess of MTSL was added to the free cysteine residues and the mixture was incubated with the protein at 4˚C overnight. After the incubations, excess MTSL was removed using a PD-10 desalting column.

The labelling efficiency, which is defined as the molar ratio of attached spin labels to cysteine residues, was found to be between 50-60% for double-cysteine mutants and approximately 90% for the single-cysteine mutant.

***Sarkosyl Extraction from Tau187 Cell Lines and AD Brains***

jR2R3 P301L fibrils were transfected in H4 cells that overexpressed the mClover3-Tau187-P301L construct, as previously described. After incubating the transfected cells overnight, the cells were washed and collected. Tau187 puncta were extracted a solution containing 2% sarkosyl in A68 buffer (10 mM Tris-HCl pH 7.5, with 10% sucrose, 0.8 M NaCl, 1mM EGTA). The A68 buffer was supplemented with PMSF and a protease inhibitor. A manual cell scraper was employed to collect the cell extracts. The samples were then sonicated for 15 seconds at 30% amplitude and incubated at 37˚ C for 30 minutes. After incubation, cell extracts were centrifuged at 113,000 g for 20 minutes at 25˚C. The sarkosyl-soluble fraction was collected and the remaining pellets were washed with 30mM Tris-HCL (pH 7.5). The samples were then centrifuged again at 113,000 g for 20 minutes at 25˚C. The sarkosyl-insoluble fractions were resuspended in 30 mM Tris-HCl (pH 7.5) and sonicated for 15 seconds at 30% amplitude.

Brain samples used in this study were sourced from the New York Brain Bank at Columbia University. The specific region of the brain from which the samples were obtained was the BA38 temporal pole, and the specimens showed neuropathologic changes associated with Alzheimer's disease. Before processing, the samples were equilibrated to -20˚C for 30 minutes, and then sliced into 0.6 micron-wide sections to accumulate a total of 0.5 grams of material. The brain tissue was homogenized in 2% sarkosyl in A68 buffer (10 mM Tris-HCl, pH 7.5, containing 10% sucrose, 0.8 M NaCl, 1 mM EGTA). At 6% (w/v) using a glass Dounce homogenizer. PMSF and protease inhibitors were added to the A68 buffer. After homogenization, the samples were centrifuged at 113,000 g for 20 minutes at 25˚C. The sarkosyl-soluble fraction was collected, and the pellets were washed with 30 mM Tris-HCl (pH 7.5). Samples were centrifuged again at 113,000 g for 20 minutes at 25˚C. The sarkosyl-insoluble fractions were resuspended in 30 mM Tris-HCl (pH 7.5) and sonicated for 15 seconds at 30% amplitude.

***Western Blotting***

The protein concentrations of the samples were determined using a Pierce BCA Protein Assay Kit from Thermo Fisher Scientific. Standard SDS-PAGE was performed using both the sarkosyl-soluble and sarkosyl-insoluble fractions derived from cell and AD brain extractions. The samples were loaded on a 4–20% Mini-PROTEAN® TGX™ Precast Protein Gel from Bio-Rad. Precision

Plus Protein Dual Color Standards ladder from Bio-Rad was used as a reference for molecular weights. For the western blot staining, the MC1 antibody hybridoma conditioned media was employed, which was kindly provided by the lab of Peter Davies. The immunoblots were imaged using a Licor Gel Imaging Station and subsequently analyzed with Fiji imaging software.

***DEER***

To prepare the heparin fibrils for the DEER analysis, a sample mixture was created by mixing tau187(spin-labeled) and tau-cys-less at a 1:20 molar ratio (5 μM:100 μM) molar ratio and incubating it with heparin at a 1:4 molar ratio of tau to heparin. The mixture was incubated at 37°C for 24 hours. For preparing jR2R3 P301L seeded tau fibrils, instead of heparin, 50% seeds (by molar percentage) were added to the same mixture and incubated at 37°C for 24 hours. The formation of fibrils was verified using ThT binding assay and TEM analysis. After fibrillization, the fibrils were centrifuged at 15000 rpm to pellet them, and the supernatant was discarded. The pellets were washed three to four times with $D_2O$ based 20 mM HEPES buffer to eliminate any remaining monomers. The final pellets were resuspended in 28 μL of $D_2O$ based 20 mM HEPES buffer, mixed with 12 μL D8-glycerol (making up 30% of the total volume), and transferred to a 2 mm diameter quartz tube before being flash-frozen using liquid nitrogen.

The four-pulse DEER experiments were conducted at 65 K on a Q-band Bruker E580 Elexsys pulse EPR spectrometer operating at approximately 34 GHz, and equipped with a 300 W Traveling Wave Tube (TWT) amplifier. The DEER pulse sequence used was: $p_{obs}/2 – t_1 – p_{obs} – (t-p_{pump}) – (t_2-t) – p_{obs} – t_2 – echo.$ Rectangular observe pulses were used with lengths set to $p_{obs}/2$ = 10-12 ns and $p_{obs}$ = 20-24 ns. A chirp πpump pulse was applied with a length of 20-24 ns and a frequency width of 133 MHz. The observed frequency was set 150 MHz higher than the center of the pump frequency range. $t_1$ was 180 ns and $t_2$ was varied between 1.8 ms and 2.4 ms. The DEER experiment was accumulated over ~approximately 12 hours.

The DEER data analysis was carried out using LongDistances software with Tikhonov regularization. The background signal, which arises from non-specific spin-spin interactions other than the intramolecular spin pairs and can interfere with the data, was minimized by using spin dilution with non-spin labeled tau. Additionally, singly spin-labeled tau (at site 351) was used to prepare fibrils, and the resulting signal was fit to determine the background dimensionality, which was found to be approximately 2 for the fibrils. These values were used for background subtraction for the respective fibrils. Other parameters were kept constant for all data to enable accurate comparison between samples. The background-corrected signal was transformed to a distance distribution, $P(Ree)$, by Tikhonov regularization with non-negativity constraint employing LongDistances software by Altenbach. The error in $P(Ree)$ was estimated using the bootstrapping method available in the LongDistances software.

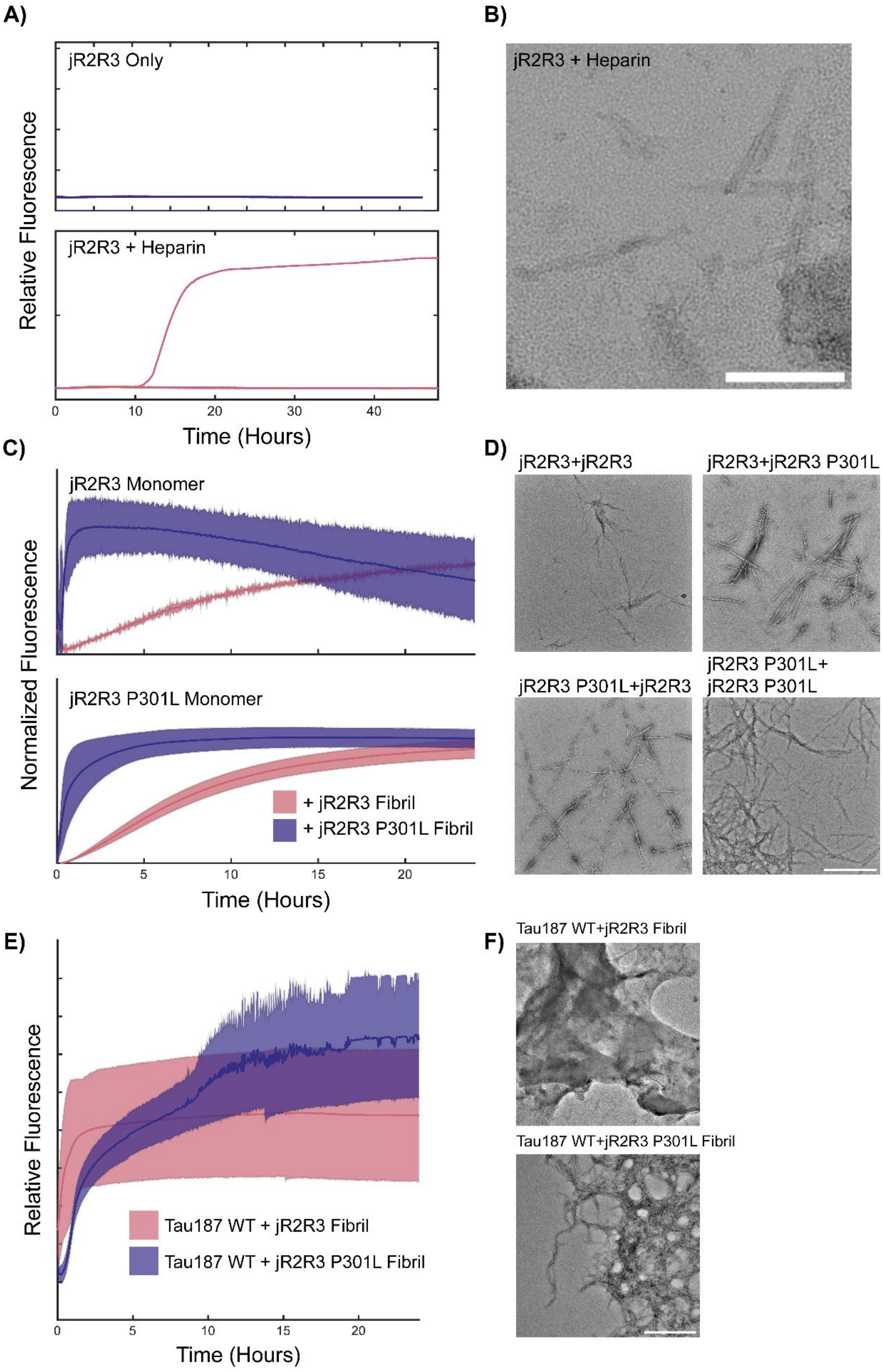
A)
jR2R3 Only
jR2R3 + Heparin
Relative Fluorescence
0
10
20
30
40
Time (Hours)
B)
jR2R3 + Heparin
C)
jR2R3 Monomer
jR2R3 P301L Monomer
Normalized Fluorescence
+ jR2R3 Fibril
+ jR2R3 P301L Fibril
0
5
10
15
20
Time (Hours)
D)
jR2R3+jR2R3
jR2R3+jR2R3 P301L
jR2R3 P301L+jR2R3
jR2R3 P301L+
jR2R3 P301L
E)
Relative Fluorescence
Tau187 WT + jR2R3 Fibril
Tau187 WT + jR2R3 P301L Fibril
0
5
10
15
20
Time (Hours)
F)
Tau187 WT+jR2R3 Fibril
Tau187 WT+jR2R3 P301L Fibril

**Fig. S1. Monitoring Aggregation of Tau Constructs by ThT and TEM** A) Among five jR2R3 reactions, only one demonstrated aggregation over a 48-hour period post-heparin addition (rose), as indicated by ThT. Isolated jR2R3 displayed no observable aggregation (purple). B) TEM imaging of heparin-seeded jR2R3fibrils revealed short, fragmented fibrillar structures. Scale bar: 200 nm. C) Both jR2R3 (rose) and jR2R3 P301L (purple) effectively seeded fibril formation in naïve monomers of themselves and of each other, with jR2R3 P301L promoting faster aggregation in both cases. D) TEM images of the four seeding reactions demonstrate clear fibrillar assemblies in each case. Scale bar: 500 nm. E) ThT fluorescence assays for Tau187 WT (rose) and Tau187 P301L (purple), demonstrating fibril formation when seeded by jR2R3. F) Corresponding TEM images of the seeding reactions, revealing the presence of fibrillar structures. Scale bar: 300 nm.

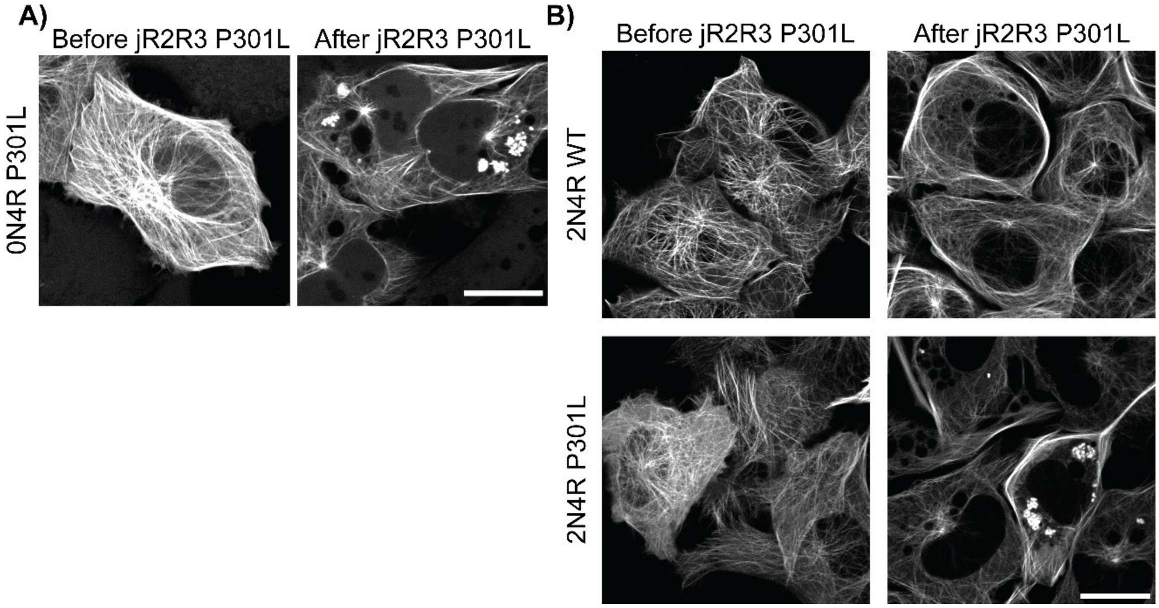


**Fig. S2. Cellular Seeding of jR2R3 and jR2R3 P301L** A) 0N4R P301L was seeded with jR2R3 P301L and puncta formation was observed after 12 hours. Puncta were observed in 13 ± 6% of cells. Scale bar: 20 µm. B) 2N4R WT and 2N4R P301L were seeded with jR2R3 P301L and puncta formation was observed after 12 hours. 2N4R had the least observed cells with at least one punctum, with 2N4R WT having 2 ± 1.7% and 2N4R P301L having 12 ± 2.4%. Scale bar: 20 µm.

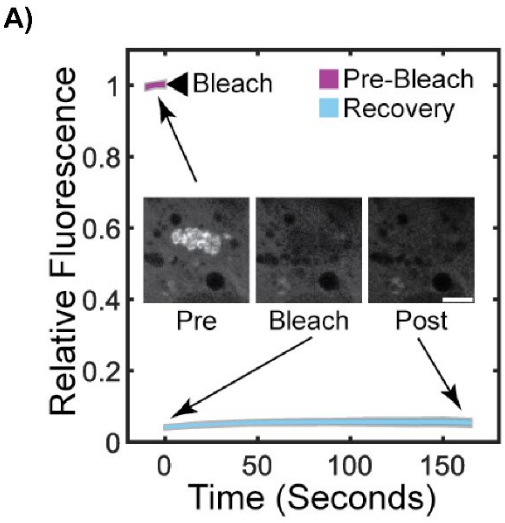


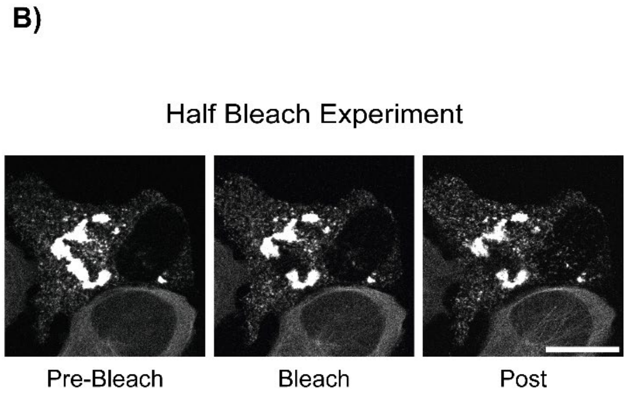


**Fig. S3. FRAP Analysis of Tau Assemblies** A) FRAP confirmed the presence of a highly immobile tau population within the puncta structures. Scale bar: 15 µm. B) Larger tau assemblies were partially subjected to FRAP, revealing no observable recovery. Scale bar: 15 µm.

A)

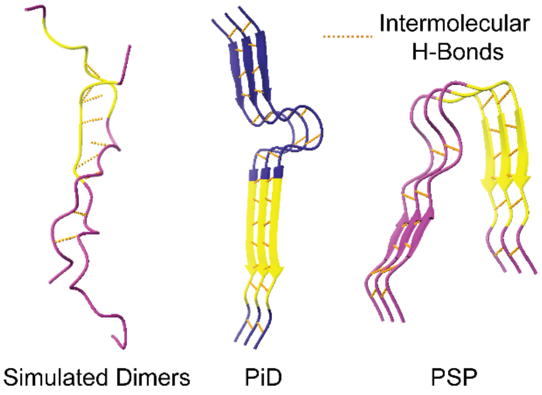


B)

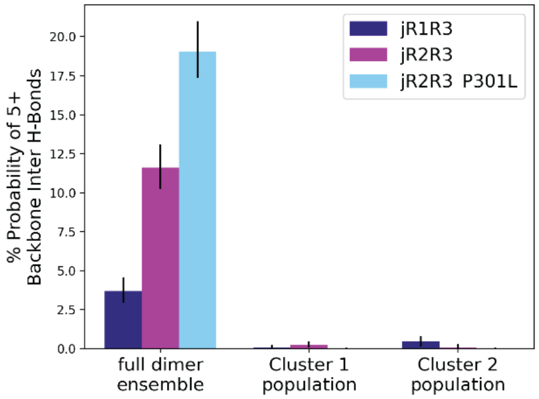


**Fig. S4. Dimer Simulations Show Increased Intermolecular Hydrogen Bonding** A) Representative conformations of dimer, with intermolecular hydrogen bonds depicted as yellow dots. A dimer conformation with both monomers in open states and featuring nine intermolecular backbone hydrogen bonds is shown on the left, compared to the structures from PiD (middle) and PSP (Right). B) Analysis of dimer simulations for jR2R3, jR2R3 P301L, and jR1R3, showing differences in intermolecular hydrogen bonding. jR1R3 exhibits reduced intermolecular hydrogen bonding compared to jR2R3 and jR2R3 P301L. Bars represent 90% confidence intervals. Oligomers with high hydrogen bonding (i.e., 5 or more intermolecular backbone hydrogen bonds) are rarely formed when one monomer is in a closed conformation.

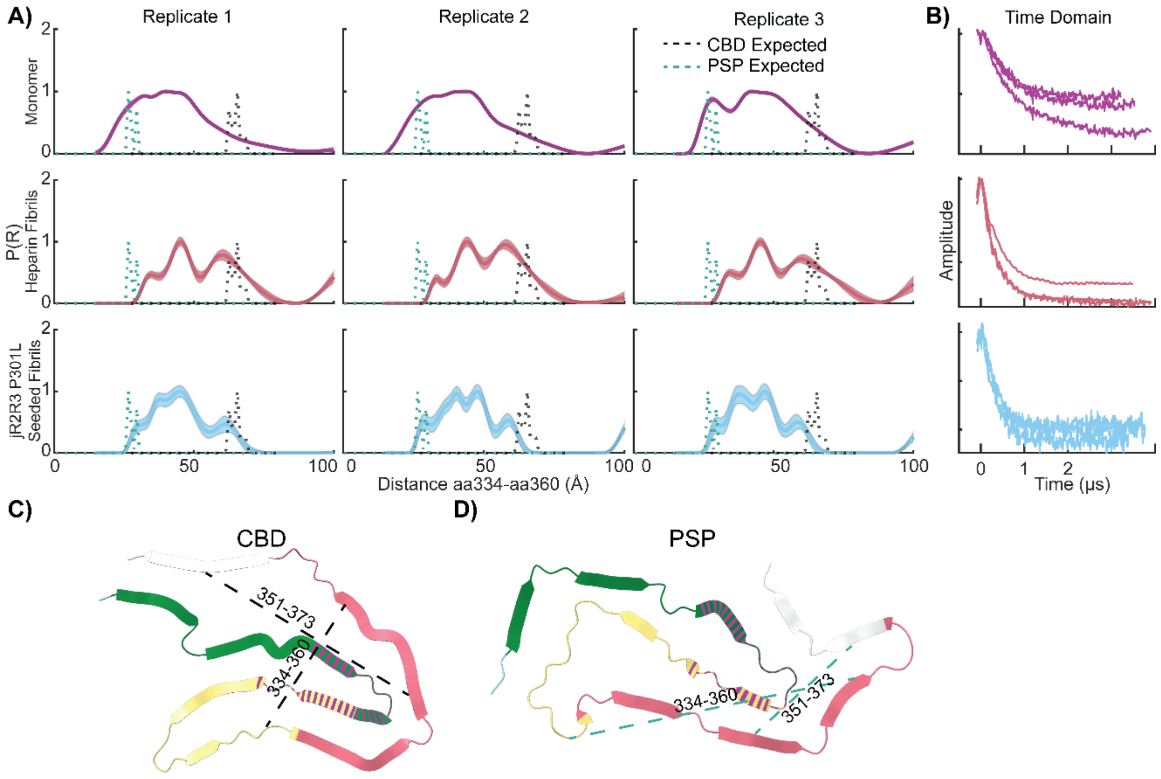


**Fig. S5. Comparative Analysis of DEER Measurements for Monomeric and Fibrillar Tau Configurations aa334-aa360 Configurations** A) DEER distance distribution plots (P(R)) for the 334-360 spin-pair distances in various tau configurations. The monomer is represented in purple, jR2R3 P301L seeded fibril in cyan, heparin fibril in rose, while the predicted distances for CBD are in black and PSP are in teal. Triplicates of each condition are plotted across the columns. B) The time domain data for each of the replicates plotted in (A). C) Hypothetical representation of distances expected if a pure CBD fold was sampled. D) Hypothetical representation of distances expected if a pure PSP fold was sampled. The figure provides insight into the conformational distinctions between tau monomers, jR2R3 P301L-seeded fibrils, and pathological tau configurations.

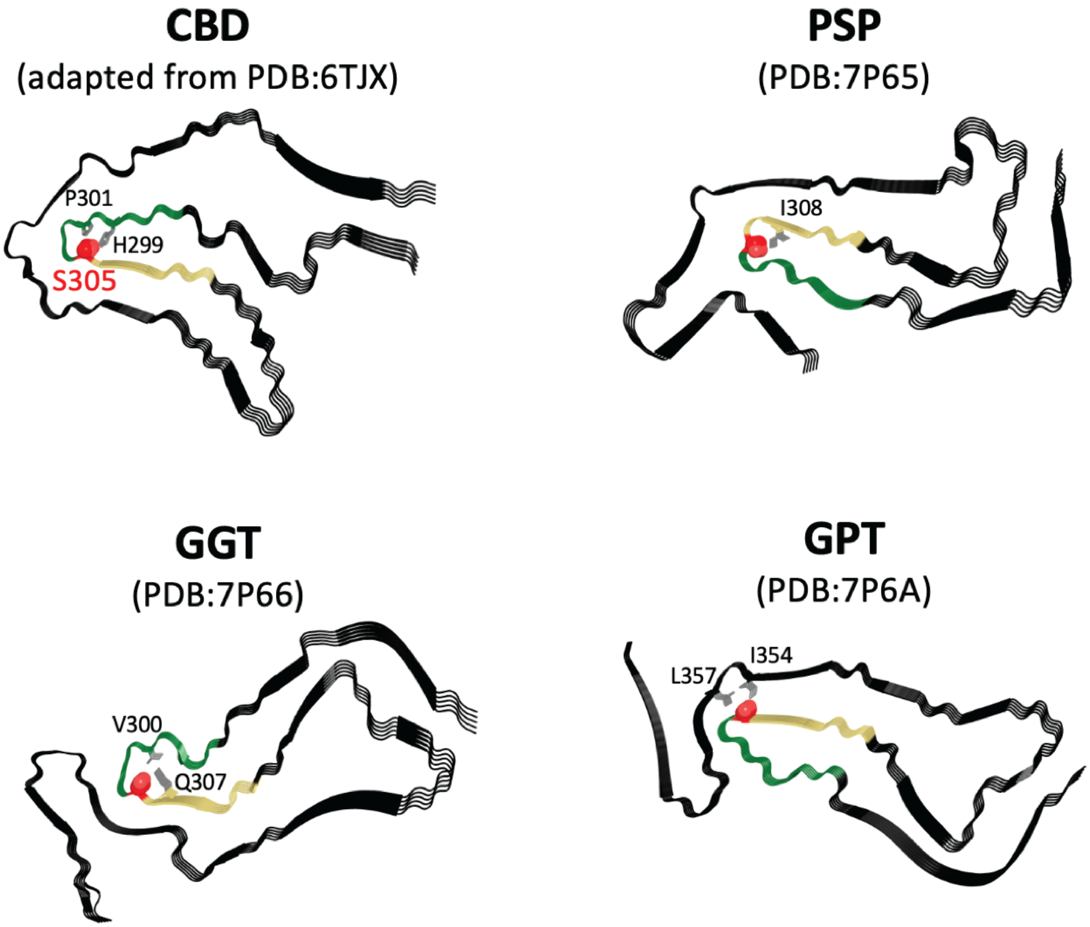


**Fig. S6. Orientation of S305 in 4R Tauopathies** The relative orientation of S305 (represented as red spheres) in CBD, PSP, GGT, GPT is illustrated. The region of the R2 repeat encompassed by jR2R3 is colored in green and that of R3 is colored in sand. In CBD, PSP, and GGT the S305 is oriented inwards towards the rest of the strand loop strand sequence while in GPT it points outwards and is stabilized by the second layer of the fibril. Residues closely interacting are shown as grey sticks.
